# Optimised whole-genome CRISPR interference screens identify ARID1A-dependent growth regulators in human induced pluripotent stem cells

**DOI:** 10.1101/2022.10.03.510590

**Authors:** Sunay Usluer, Pille Hallast, Luca Crepaldi, Yan Zhou, Katie Urgo, Cansu Dincer, Jing Su, Guillaume Noell, Kaur Alasoo, Omar El Garwany, Sebastian Gerety, Ben Newman, Oliver M. Dovey, Leopold Parts

**Affiliations:** Wellcome Sanger Institute, Cambridge, United Kingdom; Department of Computer Science, University of Tartu, Tartu, Estonia

## Abstract

Perturbation of gene function is a powerful way to understand the role of individual genes in cellular systems. Whole-genome CRISPR/Cas-based screens have parallelized this approach and identified genes that modulate growth in many contexts. However, the DNA break-induced stress upon Cas9 action limits the efficacy of these screens in important models, such as human induced pluripotent stem cells (iPSCs). Silencing with a catalytically inactive Cas9 is a less stressful alternative, but has been considered less effective so far. Here, we first tested the efficiency of several dCas9 fusion proteins for target repression in human iPSCs, and identified dCas9-KRAB-MeCP2 as the most potent. We then produced monoclonal and polyclonal cell lines carrying this construct from multiple iPSC donors, and optimized genome-wide screens with them. We found silencing in a 200bp window around the transcription start site to be as effective as using wild-type Cas9 for identifying essential genes in iPSCs, but with a reduced cost due to better cell survival. Monoclonal lines performed better, but data from polyclonal lines were of sufficient quality for screening for larger effects. Finally, we performed whole-genome screens to identify dosage sensitivities that depend on the functionality of ARID1A, a commonly mutated cancer and developmental disorder gene. We observed differential growth upon depletion of NF2, TAF6L, EZH2, and PSMB2 genes in ARID1A+/-lines compared to wild type, and an enrichment of proteasome genes. Further, we confirmed that the context-specific growth decrease was phenocopied by treating the cells with a proteasome inhibitor, suggesting a pharmacologically targetable synthetic lethal interaction between the proteasome and ARID1A. We propose that many more plausible targets in challenging cell models can be efficiently identified with our approach.

## INTRODUCTION

Cell growth has to strike a balance between supporting development and survival across the lifespan against avoiding dysregulated proliferation. The genes required for this control have been mapped using genome-scale perturbation screens in different conditions and genetic backgrounds (Miles et al., 2016), and understanding them is important for restricting tumors as well as mitigating developmental disorders (Evers et al., 2016). There is an emerging consensus of the nature of gene essentiality from screens in many human cell lines, with a set of established core essential genes that are required for survival in most tested contexts, as well as a set of context-dependent vulnerabilities (Behan et al., 2019; Hart et al., 2015, 2017).

The CRISPR/Cas9 system has rapidly become the gold standard for pooled survival screens that collect this important information (Shalem et al., 2014; Wang et al., 2014). Briefly, to knock out a gene, the Cas9 protein is directed to it by a guide RNA, which results in a double-stranded DNA break that is ultimately repaired by error-prone pathways leading to small insertions and deletions that often disrupt the reading frame (Mali et al., 2013). These perturbations are relatively straightforward to parallelize, which enables efficient screening (Hanna and Doench, 2020). However, the DNA break generated by Cas9 can be toxic and trigger cell death, especially in stem cell contexts (Aguirre et al., 2016; Haapaniemi et al., 2018; Ihry et al., 2018; Peets et al., 2019; Rosenbluh et al., 2017). An alternative approach that does not suffer from this limitation is CRISPR inhibition (CRISPRi) (Gilbert et al., 2013; Qi et al., 2013), which employs a catalytically inactive Cas9 (dCas9) protein that is unable to cut DNA, and is usually fused to different effector domains to improve inhibition (Alerasool et al., 2020; Yeo et al., 2018). CRISPRi efficacy is known to be dependent on targeting the right transcript at the right distance to the transcription start site, but also to vary across contexts, which motivates evaluating and establishing these dependencies in the extensively used systems (Radzisheuskaya et al., 2016; Rosenbluh et al., 2017; Sanson et al., 2018; Yeo et al., 2018).

An important context to study disease mechanisms, drug targets, and development is stem cells, and in particular induced pluripotent stem cells (iPSCs) (Liu et al., 2020), (Przybyla and Gilbert, 2022). The first genome-wide CRISPR/Cas screens in these cells have already shed light on mechanisms regulating embryonic stem cell survival and growth (Ihry et al., 2019; Mair et al., 2019; Peets et al., 2019). Cell types derived from stem cells via differentiation have been used to effectively and reversibly silence endogenous genes in cardiomyocytes (Mandegar et al., 2016), as well as to measure gene essentiality in neurons (Tian et al., 2019). Edited human iPSCs were also useful to identify early events of carcinogenesis before the accumulation of secondary mutations frequently observed in cancer lines (Wang et al., 2021). However, the potential of large-scale screening in stem cells is severely hampered by their cost, as the medium is expensive, and the cell death due to standard Cas9 action leads to requirements of very large numbers of cells. To unleash this potential, cost-effective, large-scale, effective drop-out CRISPRi screens have to be established in iPSCs, which requires understanding the efficacy of effector proteins and targeting constructs in this system.

Here, we develop an effective CRISPRi screening approach in iPSCs, and use it to map mutation-specific and drug-sensitive genetic dependencies. First, we compare multiple fusion proteins, and identify dCas9-KRAB-MeCP2 as the most potent one. We then establish the characteristics of successful guide RNAs that effectively repress their target using this fusion, identifying both the activity window as well as highlighting the importance of correct transcript selection. Next, we conduct whole-genome screens to determine the coverage requirements for an effective drop-out screen in iPSCs, and compare these to standard CRISPR screens. Finally, we demonstrate the efficiency of CRISPRi screens to identify nocodazole-dependent gene essentiality, as well as genes that impact growth differently as a result of a mutation to the ARID1A cancer and developmental disorder gene.

## RESULTS

Effective gene silencing with CRISPRi depends on the fusion protein used and the choice of guide RNAs. Therefore, we first explored the potency of dCas9 alternatives in human iPSCs, and identified the effect of guide RNA features, such as proximity to different transcription start sites.

### A rapid gene silencing efficacy test identifies the dCas9-KRAB-MeCP2 fusion protein as the most potent in human iPSCs

In order to identify the most potent dCas9 fusion construct, we compared three previously reported versions: dCas9-KRAB with piggyBAC delivery, KRAB-dCas9 with lentiviral delivery, and dCas9-KRAB-MeCP2 with piggyBAC delivery (Yeo et al., 2018). We produced polyclonal and monoclonal lines in iPSCs, and the human myeloid leukemia cell line K562 for each, and measured their efficacy in a dual fluorescence reporter system (Figure 1A-1B). As expected, we observed reduced intensity of the targeted GFP gene in nearly all cases, with the most effective monoclonal lines outperforming the polyclonal ones (Figure 1C). In both K562 and iPSCs, the most successful clone was a monoclonal line with the dCas9-KRAB-MeCP2 construct, so we chose these lines for our screening applications, and retained the polyclones for reference as well.

**Figure 1.**
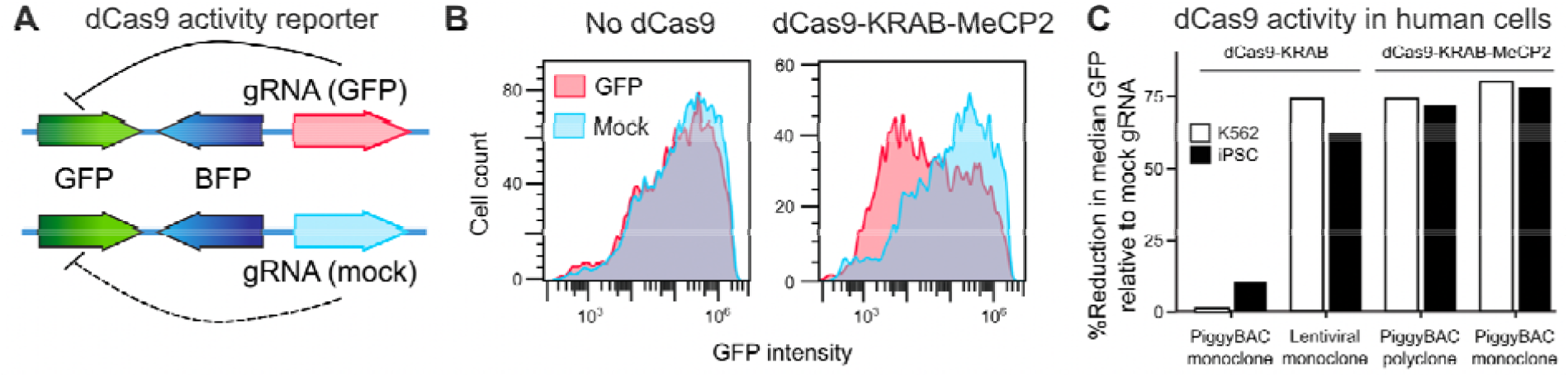
Identifying potent CRISPRi reagents in human iPSCs. **A**. Design of a dCas9 activity reporter based on internal BFP control, and GFP-targeting gRNA comparison against a mock gRNA. **B**. Frequency (y-axis) of GFP intensity (x-axis) for GFP-targeting gRNA (red) and mock control (blue) for dCas9 negative cells (left panel) and dCas9-KRAB-MeCP2 positive cells (right panel). **C**. dCas9 construct activity, quantified as the fraction of median GFP abundance with GFP-targeting gRNA compared to mock gRNA in BFP-positive cells (y-axis) for two different dCas9 constructs of different clonality and delivery method (x-axis) in K562 cells (white bars) and iPSCs (black bars).

### dCas9-KRAB-MeCP2 silencing window can extend to 1.4 kb

After picking the fusion protein to work with, we characterised the determinants of successful gene repression with dCas9-KRAB-MeCP2 in iPSCs, starting with transcription start site (TSS) distance. To determine the optimal target window, we designed a diverse gRNA library against various non-coding regions (including alternative TSSs) and coding sequences (Supplementary Figure 1).

We first targeted all NGG protospacer-adjacent motifs (PAMs) in 100 base pairs downstream of the TSS for 252 essential genes. On average, downregulation was most potent in the 20-40 bp downstream window (mean log_2_ fold change=-2.5), with a small but significant decrease of efficacy elsewhere (weakest mean log_2_ fold change=-2.2, Figure 2A). We next tiled all PAMs in a broader window of -200 bp to 300 bp around the TSS for 17 essential genes. We observed the strongest downregulation in the 0 to 100 bp window just downstream of the TSS, but more tempered effects further away (mean log_2_ fold change -2.05 vs. -1.81; Figure 2B).

**Figure 2.**
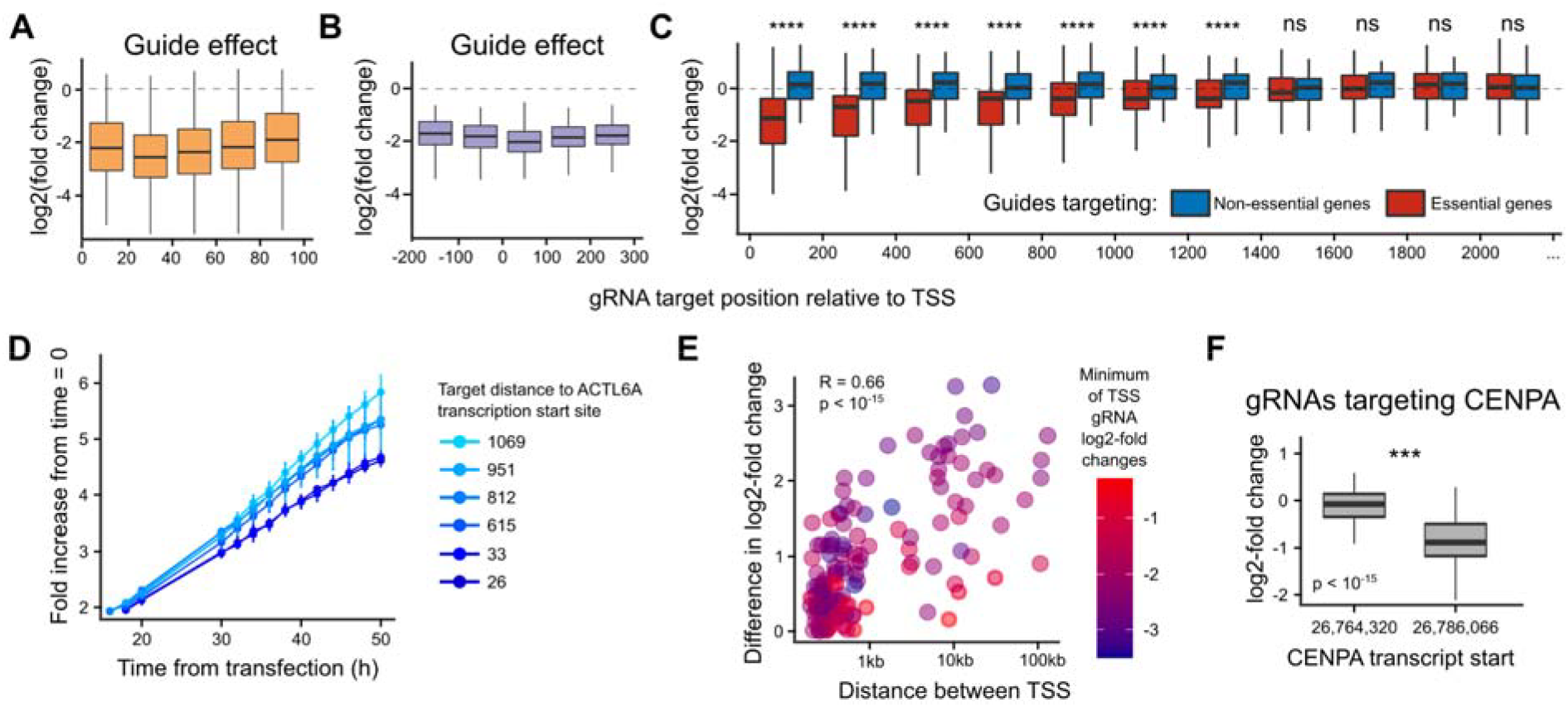
**A**. gRNA log_2_ fold change in frequency (y-axis) at different positions relative to the transcription start site (x-axis). Box: median and quartiles; whiskers: 95th percentile; dashed line: no effect. Data for 6622 gRNAs targeting 252 essential genes. **B**. As A., but data for 1816 gRNAs targeting 17 essential genes in a different range. **C**. As A., but data for 45065 gRNAs targeting 451 essential genes (red) and 4122 gRNAs targeting 49 non-essential genes (blue) in a different range. ****: Wilcoxon test p-value < 10^−5^, ns: not significant. **D**. Fold increase in confluency from time of transfection (y-axis) across time (x-axis) for gRNAs targeting ACTL6A at different distances to its transcription start site (colors). **E**. Guides targeting distant TSSs of the same gene have different effects. Difference in average log_2_-fold change of guides targeting two transcripts (y-axis) according to transcription start site separation (x-axis). Color: stronger of the two transcripts’ downregulation phenotypes. **F**. log_2_-fold change of gRNAs targeting CENPA gene (y-axis) for gRNAs targeting two different transcription start sites (x-axis). Box and whiskers: as A.

Given the relative evenness of guide efficacy in these windows close to the TSS, we finally tested even more distal targets. We re-used a design that tiled coding sequences of 440 essential genes depleted in a genome-wide screen (Supplementary Figure 2), and observed significant depletion as far as 1.4kb from the TSS, with effects further out indistinguishable from controls (Figure 2C). The decrease of efficacy was gradual, with an average log_2_-fold change for this library reducing from -1.2 within 100bp of the TSS to -0.4 at 1.2-1.4 kb away. To validate the distance-dependence of depletion phenotypes, we cloned six gRNAs targeting the essential ACTL6A gene at distances to its TSS ranging from 26bp to 1069 bp, infected a monoclonal dCas9-KRAB-MeCP2 iPSC line with them, and monitored growth under a live imaging system. We observed a decrease in growth rate that was proportional to TSS proximity (Figure 2D), confirming that the phenotypic impact can be modulated in the cell population by varying the distance of the targeting guide to the TSS.

### Targeting the correct transcription start site is important for CRISPRi efficiency

As the TSS can be cell-type dependent, picking the correct one is required for designing efficient CRISPRi reagents. To measure the impact of TSS annotation, we picked genes with two transcription start sites separated by at least 200 bp from the tiling experiment, and calculated the difference in mean log_2_ fold change of the gRNAs targeting the two transcripts. We found that the CRISPRi phenotype change depended on the proximity of the two TSSs, with longer distances leading to larger differences in phenotype (Pearson’s R=0.66, p < 10^−15^, Figure 2E). As an example, CENPA is a core essential gene with two alternative TSSs separated by 20 kb, and guides targeting the alternative TSS are substantially more depleted than guides targeting canonical TSS (median log_2_ fold change 0.0 vs -0.9, Figure 2F). Overall, TSS annotation is important for CRISPRi efficiency with the KRAB-MeCP2 construct, but, consistently with observations from tiling experiments above, only when the transcripts are separated by more than 1kb.

### CRISPRi screening in iPSCs efficiently identifies essential genes

To measure the performance of dCas9-KRAB-MeCP2 construct in identifying essential genes on the genome scale, we conducted screens in human iPSC and K562 cell lines with the Dolcetto library (Sanson et al., 2018). We aimed for 200x coverage during infection, maintained 500x coverage throughout the remainder of the screen across technical replicates, and had mean coverage of 443 for sequencing libraries (Figure 3A; Supplementary Figure 3, Methods).

**Figure 3.**
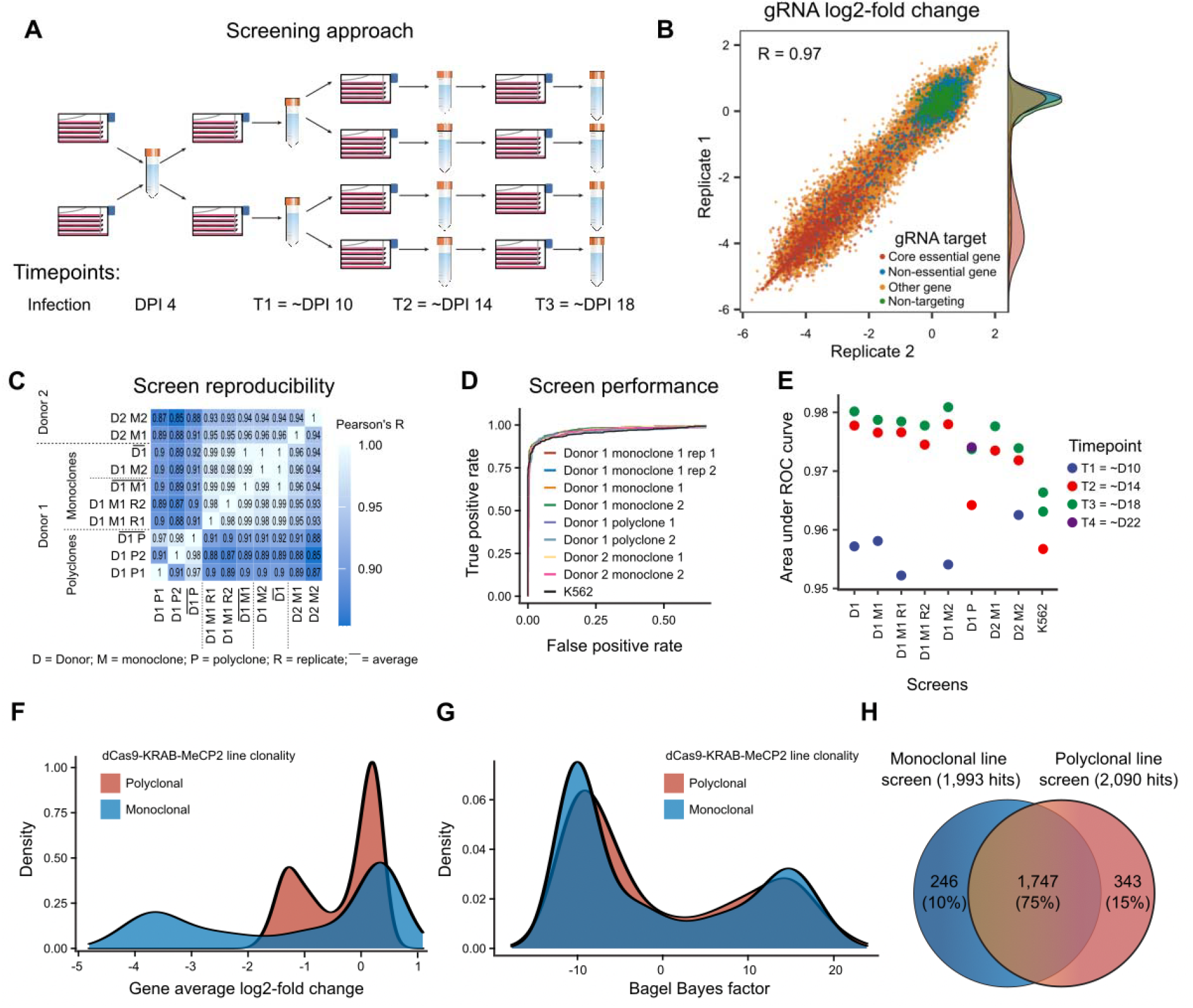
**A**. Screening approach. **B**. Reproducibility of genome-wide screens in monoclonal iPSC lines. gRNA log_2_-fold change in replicate 1 (y-axis) and replicate 2 (x-axis). Red: essential genes; blue: non-essential genes; yellow: other genes; green: non-targeting controls. **C**. Concordance. Pearson’s R of gene mean log_2_-scale gRNA fold changes (color) for screens performed in different donors, clonalities, clones, and replicates (x,y axis). **D**. Performance. True positive rate (y-axis) at different false positive rates (x-axis) for separating gold standard essential from non-essential genes in different screens (colors). **E**. Time-dependence. Area under the TPR-FPR curve (y-axis) for different screens (x-axis) across time (colors). Blue: T1 (about day 10), red: T2 (about day 14), green: T3 (about day 18), purple: T4 (about day 22). **F**. Density (y-axis) of gene average log_2_-fold changes (x-axis) of gold standard essential and non-essential genes as assayed in monoclonal (blue) and polyclonal (red) cell lines. **G**. As F., but Bayes factor computed from BAGEL on x-axis. **H**. Venn diagram of hits found with monoclonal and polyclonal line screens.

The whole genome CRISPRi drop-out screens successfully identified essential genes in both iPSCs (Figure 3) and K562 cells (Supplementary Figure 4). The screens were highly reproducible between replicates and different iPSC lines (Pearson’s R between gene average log_2_-fold changes > 0.9; Figure 3B,C), and could successfully separate gold standard core essential and non-essential genes (Hart et al., 2014), with the area under the receiver-operator curve (AUC) above 0.9 in all cases (Figure 3D,E, Supplementary Table 2). The performance at 50x coverage of a single replicate at day 18 post-infection was very close to that of a 200x screen combining four replicates (AUC=0.977 vs 0.978). There was little difference to another biological replicate (AUC=0.978 vs 0.980), as well as to combining information from multiple biological replicates (AUC=0.980), or performing a single screen at 100x (AUC=0.974). The separation of positive and negative controls increased with every passage, with the most performance gained when going from passage two at day 8 to passage three at day 14 (AUC=0.957 vs 0.977), and with little further improvement afterwards (AUC=0.980 at day 18). Altogether, 50x coverage screens in monoclonal lines measured at day 14 have near-optimal performance in iPSCs, but polyclonal lines and earlier timepoints can be used to trade quality for cost.

Producing monoclonal cell lines from a polyclonal pool requires at least one month of extra work and carries risks of additional positively selected mutations that can confound screening results. To test whether polyclonal cell populations suffice for high quality screens, we conducted experiments in the parental polyclonal iPSC line with 70% silencing activity. We observed lower reproducibility of the biological replicates of polyclones (Pearson’s R of gene mean gRNA LFC=0.91 vs. 0.99 in monoclones), as well as worse resolution at 50x coverage to separate essential and non-essential genes (median AUC=0.97 vs. 0.98 in monoclones). However, this performance is still above the range reported in large-scale resources like the Cancer Dependency Map (Behan et al., 2019), with all of the lines screened with wild-type Cas9 showing AUCs below 0.95. In addition, the log_2_-fold changes are highly correlated between mono- and polyclones (Pearson’s R=0.92), Bayesian analysis approaches can overcome some of the reduction in the signal (Figure 3F,G), and 75% of genes identified by either of the screens are common to both (Figure 3H). Therefore, depending on the desired measurement precision, the cheaper and faster polyclonal screening will be sufficient for many purposes, and especially for identifying the strongest hits.

### CRISPRi is as accurate as CRISPR in separating reference essential and non-essential genes

Next, we asked how the performance of the KRAB-MeCP2 CRISPRi construct compares with wild type Cas9 screens. We previously conducted CRISPR/Cas9 screens in the same iPSC and K562 cell lines using a double guide RNA library targeting 18,000 genes with 3 guides per gene (Peets et al., 2019). The CRISPRi screens compared favourably, with higher AUC values (0.98 for IPSC and 0.96 for K562) than CRISPR/Cas screens (0.95 for both lines; Figure 4A). The gene-level average guide RNA log_2_ fold changes were moderately correlated between CRISPR and CRISPRi screens in the same cell line (Pearson’s R=0.42), which is in line with concordances of both knock-out and repression screens in different types of stem cells (R=0.37 to 0.53, Supplementary Figure 5). Further, of the essential genes identified in one screen, about a half were also hits in the other (1,143/2,292 for CRISPR and 1,143/2,159 for CRISPRi, Figure 4F). Overall, both systems can identify essential genes, but differences in cell line context, growth conditions, the library used, screen coverage, and other experimental parameters generate substantial variation in performance and hit lists both within and between various screen types. CRISPRi screens can be cheaper in stem cells due to the lack of double-strand break toxicity, as well as give reasonable performance using polyclonal lines.

**Figure 4.**
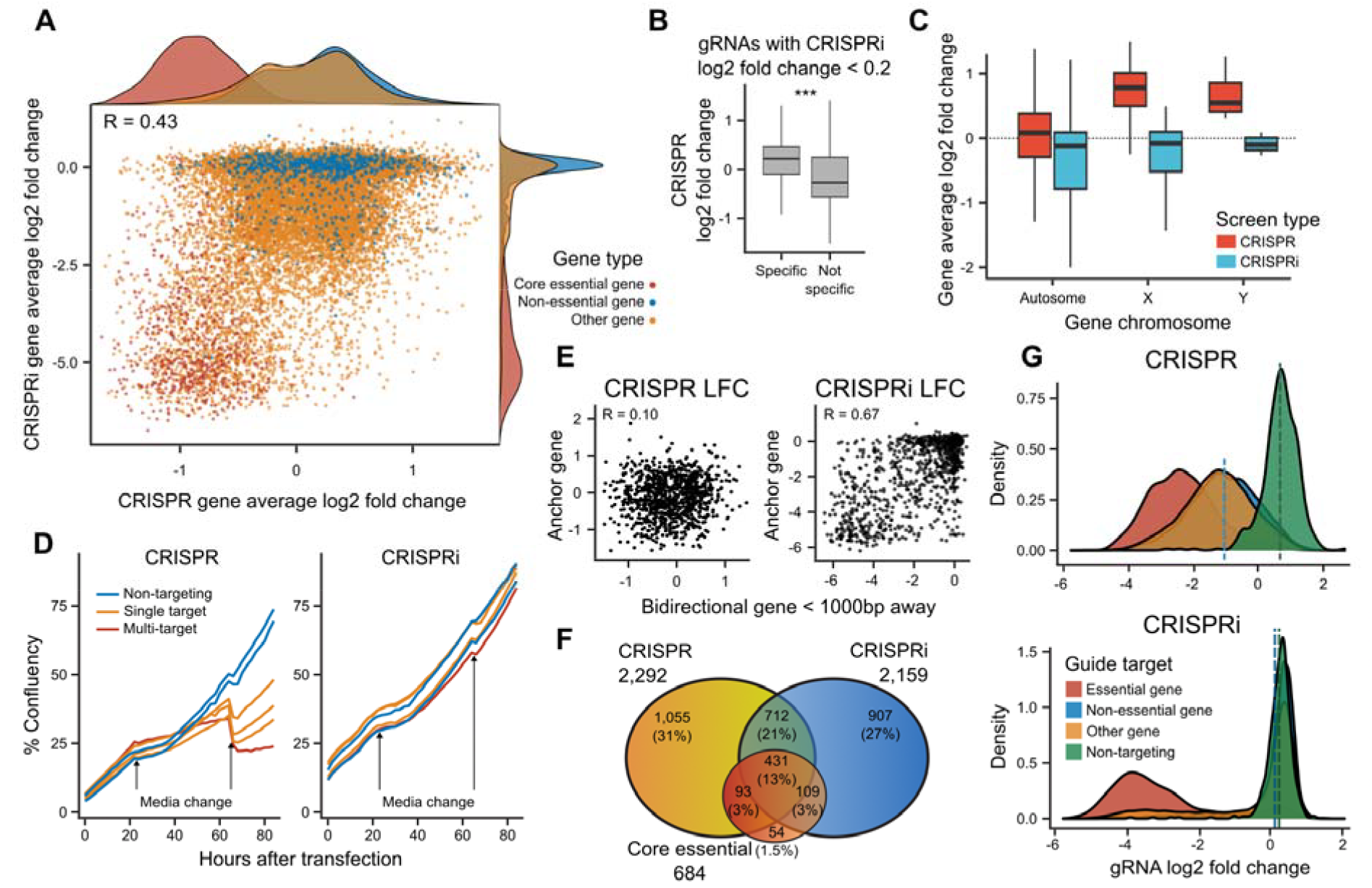
**A**. Consistency of CRISPR and CRISPRi. gRNA log_2_-fold change in CRISPRi (y-axis) and CRISPR screens (x-axis). Red: essential genes; blue: non-essential genes; yellow: other genes; green: non-targeting controls. **B**. gRNA log_2_-fold change in CRISPR screen (y-axis) for gRNAs that have no signal (absolute log_2_-fold change < 0.1) in CRISPRi screen, stratified to specific gRNAs with a single target (left), and non-specific ones (right). Box and whiskers: as in 2A. **C**. Gene average log_2_-fold change (y-axis) for genes on different types of chromosomes (x-axis) for CRISPRi (blue) and CRISPR (red) screens. Box and whiskers: as in 2A. **D**. %Confluency (y-axis) across time since transfection (x-axis) for gRNAs with different number of targets (colors). Blue: 0 targets; yellow: 1 target; red: 2 or more targets. Arrows: media change times. **E**. log_2_-fold change of gRNA (markeres) at an anchor gene (x-axis) and another gRNA at a bidirectional gene at most 1kb away (y-axis) for CRISPR screens (left panel) and CRISPRi screens (right panel). **F**. Venn diagram of overlap of CRISPR screen hits, CRISPRi screen hits, and gold standard essential genes. **G**. Density (y-axis) of gRNA log_2_-fold change (x-axis) for gRNAs in CRISPR screen (top panel) and CRISPRi screen (bottom panel). Red: essential gene targeting; blue: non-essential gene targeting; yellow: other gene targeting; green: non-targeting controls.

CRISPR and CRISPRi screens employ different mechanisms of action to perturb the target, which can bias the identified hits. For example, targeting the X chromosome with wtCas9 in lines from male donors creates DNA breaks on only one chromosome and is less toxic to cells. Thus guides targeting single copy chromosomes are enriched in CRISPR screens, but not CRISPRi screens (Figure 4B), while non-specific guides are more depleted (Figure 4C). On the other hand, the wider “blast radius” of CRISPRi can cause false-positive results (Rosenbluh et al., 2017). Indeed, when we compared nearby gene pairs with at most 1kb distance between their transcription start sites, we observed a high correlation of signal in CRISPRi screens (R=0.61), but not in CRISPR screens (Figure 4D), indicating a likely confounding due to targeting nearby TSSs.

The main drawback of CRISPR screens is double-strand break-induced cell death, which can be overcome by silencing *TP53* and p53 pathway genes (Haapaniemi et al., 2018). We first tested whether targeting *TP53* by CRISPRi in iPSCs also provides a growth advantage, and found it to confer a positive effect, but not as large as in CRISPR screens (log_2_ fold change upon TP53 perturbation 1.04 by CRISPR vs. 0.55 by CRISPRi; Supplementary Figure 6). Next, we confirmed the deleterious effect of double-strand breaks in CRISPR, but not CRISPRi screens. Indeed, the average depletion of non-targeting control guide RNAs was close to that of gRNAs targeting non-essential genes for CRISPRi screen (0.2 vs 0.1, Figure 4G), but not CRISPR screen (0.68 vs - 1.02). This implies that targeting alone with a wtCas9 construct carries a phenotypic impact in human iPSCs. Finally, we validated that this effect is dose-dependent, by comparing the growth of Cas9 and dCas9-KRAB-MeCP2 lines with guides that target zero, one, or multiple targets in the genome, and monitoring cell growth in a live imaging system. All three guides showed the same growth pattern in iPSC-dCas9-KRAB-MeCP2 lines, while the cutting at an additional number of targets caused increased cell death in iPSC-Cas9 cells (Figure 4D). Altogether, CRISPRi screens do not suffer from the dose-dependent double-strand break-induced cell death observed in CRISPR screens, but repressing the TP53 tumor suppressor still provides a moderate selective advantage.

### CRISPRi identifies context-dependent essential genes

An important screening paradigm is identification of context-specific vulnerabilities for therapeutics and diagnostics. We therefore explored the utility of CRISPRi screens for mapping genome-scale drug-gene and allele-gene interactions. We first conducted pooled CRISPRi screens to identify genes that lead to Nocodazole-dependent cell depletion (Methods). Nocodazole is an anti-mitotic agent that binds and destabilises microtubules, and also causes detachment of cells from the flask surface. The two genes most enriched in screens under Nocodazole treatment compared to standard medium were TRIP13 and BRD9 (Figure 5A). TRIP13 is a gene implicated in cell cycle control mechanisms, and its depletion is known to slow cell division, but to also allow cells to escape Nocodazole-induced cell cycle arrest (Ma and Poon, 2016, Marks et al, 2017). BRD9, however, has a TSS only 80bp apart from TRIP13, suggesting that the BRD9 signal may be an unintended effect of the CRISPRi system, and indeed, BRD9 was only a hit using CRISPRi (LFC < -3), but not CRISPR screening (LFC=0.16). Thus, the Nocodazole-dependent signal from BRD9 is a likely false positive because of its transcription start site proximity to the known TRIP13 causal gene. Therefore, CRISPRi screening can identify gene-drug interactions, but the outputs need to be carefully analyzed to avoid modes of false positives not observed in CRISPR screens.

**Figure 5.**
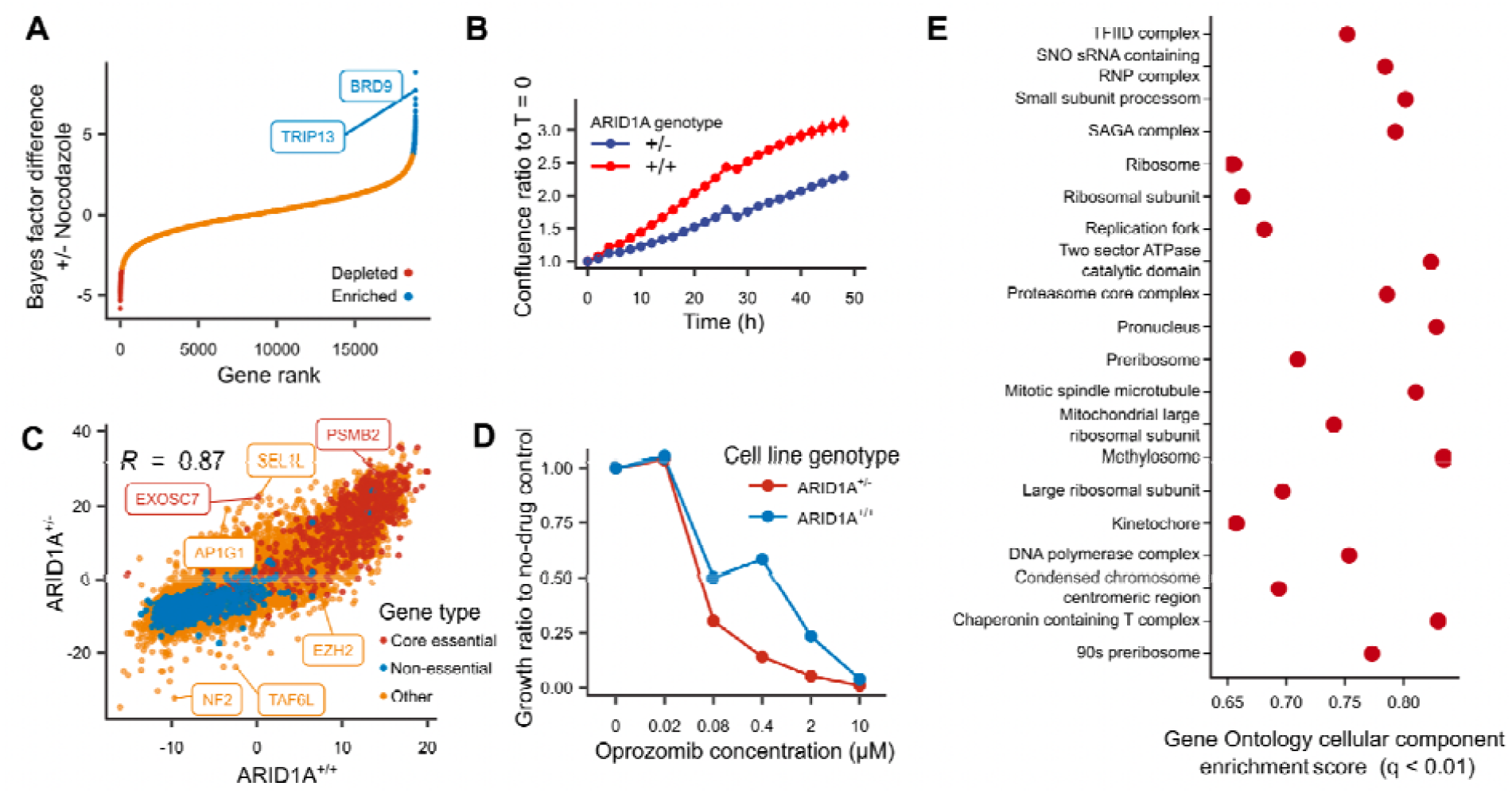
**A**. BAGEL Bayes factor difference between screens with and without Nocodazole treatment (y-axis) in increasing value (x-axis). Red: targeted genes significantly depleted in Nocodazole; blue: targeted genes significantly enriched in Nocodazole. **B**. Cell density relative to start time (y-axis) across time (x-axis) for iPSC line with wild type ARID1A genotype (red) and heterozygous loss-of-function (blue). **C**. BAGEL Bayes factor for genes (markers) for screens in ARID1A heterozygous knock-out cell line (y-axis) and no-mutation control (x-axis). Red: essential genes; blue: non-essential genes; yellow: other genes. **D**. Growth relative to no drug control (y-axis) at different concentrations of Oprozimib (x-axis) in ARID1A^+/+^ (blue) and ARID1A^+/-^ (red) cell lines. **E**. Enrichment score (x-axis) for different Gene Ontology cellular components (y-axis).

To identify mutation-specific gene effects, we screened for ARID1A-dependent vulnerabilities in iPSCs. The ARID1A gene produces a subunit of the ATP-dependent histone deacetylase SWI/SNF complex that has been implicated in developmental disorders, as well as 20% of all cancers (Pagliaroli and Trizzino, 2021), and is therefore potentially an important synthetic lethal gene dependency target. We produced isogenic iPSC lines by targeting the ARID1A coding region with transiently expressed Cas9, and picking one clonal line with a 7-nucleotide heterozygous deletion (c.2389-2395: TCCAGCAGC>TC, p. 667-669: SSS/X), and one control line with wild type sequence. The heterozygous mutation decreased ARID1A expression by 60% and resulted in slower growth compared to the wild type line (Figure 5B). To prepare the cells for CRISPRi screening, we transfected the two lines with the dCas9-KRAB-MeCP2 construct and selected monoclonal cell lines with at least 65% silencing activity in the reporter assay. We proceeded with genome-wide screens in mutant and wild-type backgrounds using the Dolcetto library as described above. The depletion signals of the two screens were less correlated compared to replicates in wild-type cells (Pearson’s R of Bagel Bayes Factors = 0.87 vs 0.94; Figure 5C), suggesting that the ARID1A mutation altered the gene essentiality profile.

We next aimed to understand the differentially depletion-sensitive genes. NF2 and TAF6L genes had the strongest increase of statistical signal in the mutant line (Methods), while AP1G1, SEL1L, and EXOSC7 had the most decrease. NF2 is a tumor suppressor frequently observed in schwannomas and shown to inhibit YAP/TAZ driven tumorigenesis (Petrilli and Fernández-Valle, 2016). The ARID1A/NF2 interaction has been observed in mouse liver cells, where the upregulation of YAP/TAZ complex by downregulation of NF2 is not enough to trigger neoplasia, but the double silencing of NF2 and ARID1A induces hepatocellular carcinoma (Chang et al., 2018). The second-strongest differential growth signal was observed upon silencing of TAF6L gene. TAF6L is a member of SAGA chromatin modifying complex, which has antagonistic roles on chromatin accessibility (Cheon et al., 2020). Targeted CRISPR screening in mouse embryonic stem cells with guide RNAs targeting epigenetic regulators found TAF6L to be important for maintaining stem cell identity in c-Myc dependent manner (Seruggia et al., 2019).

The AP1G1 gene codes for the gamma-1 subunit of adaptor-related protein complex 1, involved in the formation of clathrin-coated pits and intracellular trafficking. Mutations in AP1G1 are implicated in neurodevelopmental disorders (Usmani et al., 2021) and together with other members of SWI/SNF complex, it is reported as a proviral gene for SARS-CoV-2 infection in Calu-3 cells, as well as Vero E6 cells (Goujon et al., 2021). The second most depleted gene in ARID1A mutant lines compared to wild type was SEL1L. This gene encodes the adaptor subunit of ERAD ubiquitin ligase, which extracts misfolded proteins from the endoplasmic reticulum into the cytosol to be degraded by the proteasome (Hwang and Qi, 2018). Consistent with the role of protein degradation, inhibition of Proteasome 20S Subunit Beta 2 (PSMB2) gene, a potential drug target, resulted in more cell death in the ARID1A +/-line (Figure 5B). This sensitivity suggests a synthetic lethal vulnerability that could be therapeutically exploited. To except this differential growth effect of proteasome inhibition on ARID1A wild type and mutant lines, we applied physiological concentrations of oprozomib, a second generation proteasome inhibitor in trial for the treatment of haematological cancers (Sherman and Li, 2020), and measured the difference in growth rate. We found that ARID1A mutant line decreased in cell confluence more than wild type cells upon oprozomib treatment when compared to the no drug control (Figure 5D).

More globally, gene set enrichment analysis of all the genes ranked by the difference between the wild type and mutant line screen depletion identified the proteasome core complex among the top 20 enriched pathways (Figure 5E), as well as other pathways related to ARID1A function, like centromere complex assembly and kinetochore organization, confirming previous results (Caumanns et al., 2018; Mathur, 2018). Further, a String database (Szklarczyk et al., 2021) analysis with 200 most depleted genes identified three clusters: gene expression regulation, cell cycle regulation, and cholesterol biosynthesis regulation (Supplementary Figure 7) around ARID1A and two other members of SWI/SNF family (SS18 and SMARCE1). The SWI/SNF complex is also important in regulation of YAP/TAZ driven cell growth, and therefore indirectly implicated in different ways for ARID1A dependent gene essentiality signals. We also tested whether essentiality changes upon ARID1A dose reduction mimic those in cancer cell lines upon damaging mutations to the ARID1A gene, but found no strong signal (<u>Supplementary Note</u>).

### Context-dependent vulnerabilities replicate with different reagents and assays

We confirmed the genotype and small molecule dependent fitness effects with alternative reagents and assays. First, to validate that the reduced dose of ARID1A, rather than any other clonal mutation leads to growth rate reduction in the ARID1A mutant line, we tested six guide RNAs targeting 17, 59, 894, 987, 1160, and 1314 nt downstream of the transcription start site of ARID1A in wild type iPSCs (Methods). Two of the guide RNAs, targeting 17 nt and 1314 nt downstream of TSS, caused substantial growth retardation in the first 36 hours after transfection (Figure 6A).

**Figure 6.**
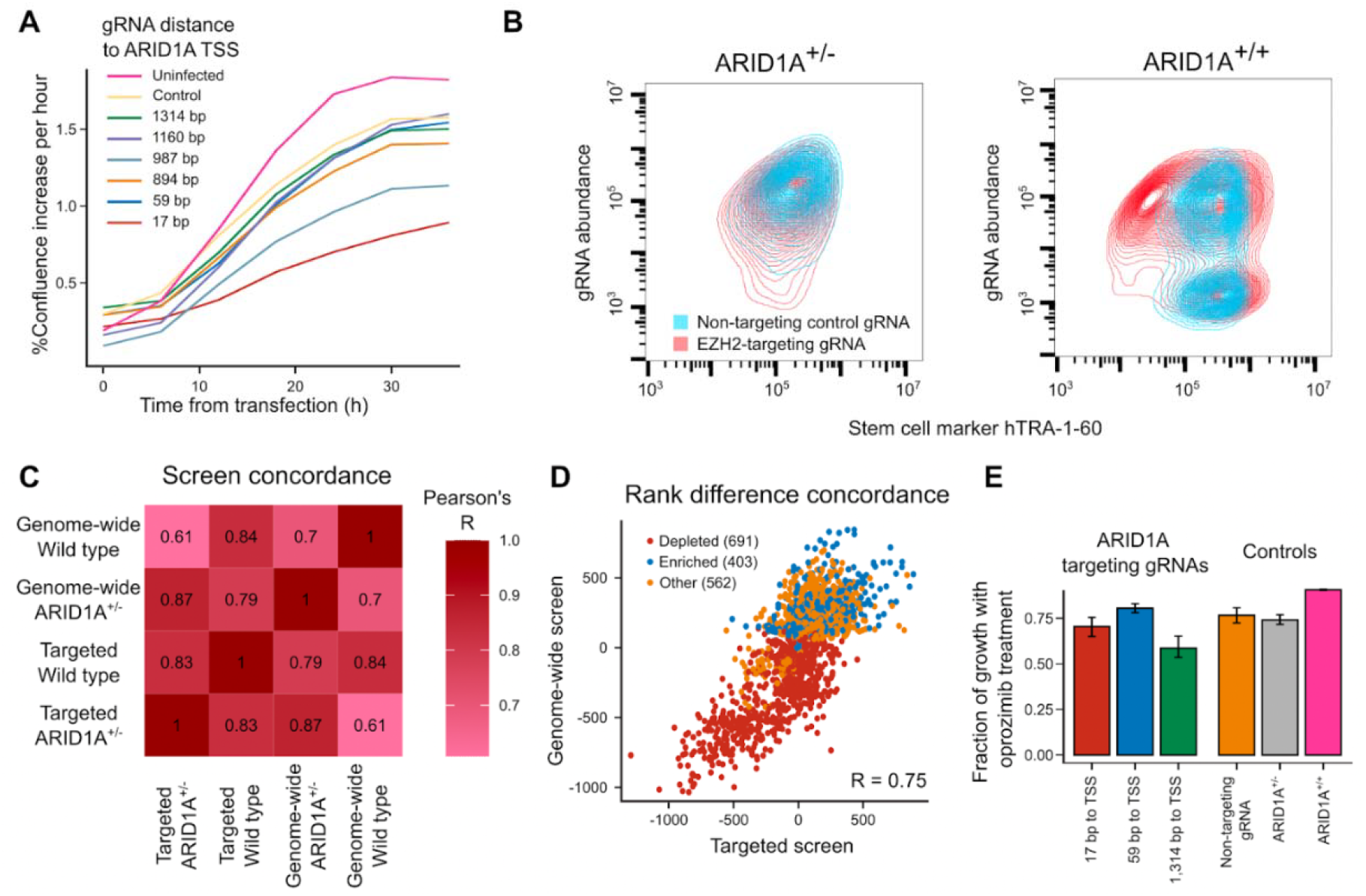
**A**. Percent confluence increase per hour (y-axis) across time from transfection (x-axis) of guide RNAs targeting different distances to the ARID1A gene transcription start site (colors) into iPSCs. **B**. Density curves of stem cell marker hTRA-1-60 signal (x-axis) for different gRNA plasmid abundance proxies (y-axis) in ARID1A^+/-^ (left panel) and ARID1A^+/+^ (right panel) cell lines. Red: EZH2-targeting gRNA; blue: non-targeting control gRNA. **C**. Screen concordance (Pearson’s R of gene average log_2_-fold changes; colors) of genome-wide and targeted CRISPRi screens in ARID1A+/+ and ARID1A+/-cells (x and y axes). **D**. Rank difference of genes (markers) between CRISPRi screens in ARID1A+/+ and ARID1A+/-cells in a genome-wide screen (y-axis) and targeted followup screen (x-axis). Red: depleted genes in the genome-wide screen; blue: enriched genes; orange: other genes. **E**. Fraction of growth upon oprozomib treatment compared to no drug control (y-axis) for ARID1A targeting gRNAs (x-axis; first three bars) and controls (last three bars).

Second, we confirmed the impact of silencing EZH2, coding for the catalytic domain of histone methyltransferase Polycomb Repressive Complex 2 (Figure 5B), which was more essential in the wild type screen compared to the one in ARID1A mutant line. We transfected cells with either EZH2 targeting or non-targeting guideRNAs, and after nine days in culture, observed substantial morphology changes upon targeting compared to non-targeting guides in ARID1A+/+ cells and EZH2 silenced ARID1A+/-cells (Supplementary Figure 8). Further, 32% of EZH2 silenced ARID1A+/+ cells were negative for the stem cell marker hTRA-1-60 (Figure 6B), indicating likely differentiation from stem cell state towards a less propagating lineage with different morphology and expression profile, and implicating ARID1A as a causal intermediate factor in this process.

Third, we validated hits from the whole-genome screens. We compared the signal between CRISPRi screens in ARID1A+/+ and ARID1A+/-lines and picked 1657 genes with large differences (Methods) to re-measure in a followup experiment. Screens in the wild type and mutant lines using a gRNA library against these genes were consistent with the original ones both in absolute terms (Pearson’s R of gene LFCs > 0.84, Figure 6C and Supplementary Figure 9), as well as for difference in the mutant line (Pearson’s R of rank changes 0.75, Figure 6D).

Finally, to confirm that sensitivity to proteasome inhibition is due to ARID1A gene targeting, we treated the ARID1A wild-type lines with the two strongly growth retarding ARID1A-targeting gRNAs, one weak gRNA, a negative control gRNA, and no gRNA, as well as with oprozomib (1μg/ml) for one day and compared growth to a non-treated controls, and the ARID1A +/-line. The vulnerability of hiIPSC cells to oprozomib treatment was dependent on ARID1A targeting, with gRNAs leading to stronger growth effect in human iPSCs also giving rise to stronger phenotypes upon proteasome inhibition, resembling the growth of the mutant line (Figure 6E). Thus, we have demonstrated that an important developmental disorder and cancer gene can potentially be therapeutically targeted with a small molecule in a dose-dependent manner.

## DISCUSSION

We presented the first systematic investigation of genome-wide CRISPR-inhibition screens in human-induced pluripotent stem cells and demonstrated that CRISPR inhibition is an efficient and safe alternative to standard CRISPR/Cas-based screening. We identified dCas9-KRAB-MeCP2 as the most potent fusion protein for target silencing in human iPSCs, with a range that can extend to 1.4kb into the coding region. Whole-genome CRISPRi screens using this construct performed as well as different novel and previously published screens with standard Cas9. We identified actionable drug sensitivity for ARID1A mutant iPSCs, and other interactions illuminating the role of ARID1A for stem cell growth and differentiation.

The best silencing modality is not easy to choose for a new experiment. For example, dCas9-KRAB is effective when integrated into the AAVS1 locus in human iPSCs (Mandegar et al., 2016), but replicating this setup is cumbersome for screening in multiple lines. The dCas9-KRAB-MeCP2 performed even better for silencing (Yeo et al., 2018), but varied in efficiency across cell types. Via testing three constructs with different delivery methods and both monoclonal and polyclonal versions, we found that monoclonal lines expressing the dCas9-KRAB-MeCP2 fusion protein showed both best silencing activity with the reporter construct and best performance in genome-wide screens. However, producing them requires an extra month of work, so the requirement for reduced noise can be weighed against the labor needed to produce subclones given the performance goals.

Another important consideration in practice is the cost required for screening at scale. Both standard CRISPR and CRISPRi screens can identify essential genes in an experiment (Gilbert et al., 2014; Horlbeck et al., 2016; Sanson et al., 2018), and while CRISPR has previously been argued to be more sensitive (Rosenbluh et al., 2017; Sanson et al., 2018), CRISPRi outperformed CRISPR in both human iPSC and K562 cells in our hands, at similar screen coverages and screen durations. In particular, we observed that a 14-day duplicate screen with 50x coverage at infection, and 200x during passaging gives near-optimal results, greatly reducing the cost per screen compared to more standard larger designs. Combined with the flexibility of choosing monoclonal lines with high efficacy, the budget and resolution can be traded off according to demand.

The dCas9 protein has to be targeted near the transcription start site for efficient screening (Gilbert et al., 2013), but the “blast radius” varies for different fusion proteins. The best window for dCas9-KRAB activity has been established to be 0 to 100 bp after the TSS (Radzisheuskaya et al., 2016; Rosenbluh et al., 2017; Sanson et al., 2018; Yeo et al., 2018). However, these reports are specific to the KRAB fusion and explored only a few target sites. We demonstrated that the target range of dCas9-KRAB-MeCP2 in iPSCs is larger, with impacts on gene expression as far as 1.6kb after the TSS, and with the efficiency gradually decreasing with distance. We propose that this understanding could be used as a way to modulate the extent of downregulation, as has been previously done with mismatched gRNAs (Jost et al., 2020). This fine-grained control would be useful to study haploinsufficiency of target genes, and other questions that require a more precise range of the gene dose. The large impact range of dCas9-KRAB-MeCP2 also decreases the false negative rate due to alternative TSS for different cell types. We showed that false negative results rarely occur for the majority of genes for which the alternative is less than 1kb away.

One important difference between CRISPR and CRISPRi screen efficiency stems from the substantial p53-mediated DNA damage response induced by wtCas9 (Aguirre et al., 2016; Haapaniemi et al., 2018). We compared growth of Cas9 and dCas9-KRAB-MeCP2 expressing iPSCs and observed that cells die upon Cas9 targeting, and that the number of target sites increases cell loss. This imposes a risk of finding false positives in amplified regions of the genome using CRISPR screening, and methods have been developed to address this (Iorio et al., 2018; Meyers et al., 2017; Perez et al., 2022). However, as the CRISPRi screens also produce false positives in overlapping transcriptional control regions, either careful analysis methods or complementary screens need to be employed.

CRISPR screens have already yielded gene-gene interactions (Horlbeck et al., 2018; Kim et al., 2022). We screened for interactions with the ARID1A gene, implicated in both developmental disorders (OMIM: 135900), as well as cancer (Pagliaroli and Trizzino, 2021), suggesting growth control to be an important aspect of its function. Suppressing the NF2 gene in ARID1A mutant background gave iPSCs a growth advantage, and although direct interaction of ARID1A and NF2 is not known, both proteins regulate pathways inhibiting oncogenic YAP/TAZ genes, and their double mutation causes hepatocellular carcinoma in mice (Chang et al., 2018; Patel et al., 2017). Two other hits, TAF6L and EZH2, are members of the SAGA and PRC2 chromatin-modifying complexes, interacting with SWI/SNF in cell fate determination (Pagliaroli and Trizzino, 2021; Seruggia et al., 2019). Further investigation of EZH2 effect with an arrayed screen demonstrated the growth advantage in ARID1+/-background was due to inhibition of cell differentiation, which further illuminates ARID1A function in cell fate determination.

Pluripotent stem cells of different provenance and capacity are poised to expand our understanding and ability to engineer disease mutations, cell types, entire organs, and courses of development (Hanna et al., 2002; Liu et al., 2020). Precise methods for genome engineering to enable control in these systems is a crucial enabling technology for this progress. Detection of context-specific effects will be key both for accurately targeting only the diseased state, as well as to chart the inherent heterogeneity that needs to be accounted for.

## METHODS

### Cell Culture

K562-Cas9 and K562-Cas9-GFP cells were cultured in RPMI supplemented with 10% FCS, 2 mM L-glutamine, 100 U/ml penicillin, and 100 mg/ml streptomycin (will be referred to as supplemented RPMI media here after). Cells were passaged 1:20 every 4 days. Wild type hIPSC (fiaj-1 and kolf2) cell lines was obtained from HipSci platform, where its karyotype and phenotype were evaluated to be normal. hIPSC lines were cultured on Vitronectin-XF (Stemcell Technologies, 07180) coated plates and in mTeSR-Plus Media (Stemcell Technologies, 100-0276). For maintenance, they were passaged every four days as clumps using ReLeSR (Stemcell Technologies, 05872) according to the producer’s protocol. All cell lines were cultured at 37°C, 5% CO2.

### Cloning Stable dCas9 Fusion Protein Expressing Lines

Three different dCas9 fusion proteins were tested for dCas9 activity in both K562 and hiPSC lines: pLX_311-KRAB-dCas9 (Addgene 96918), pB-CAGGS-dCas9-KRAB-MeCP2 (Addgene, 110824) and pPB-dCas9-KRAB (kindly provided by Dr. Qianxin Wu). The first is a lentiviral delivery vector, whereas the latter two rely on the piggyBac transposase for genome integration.

#### Lentivirus production

Lentivirus preps were produced in 293FT cells transfected with lentiviral delivery vectors together with second generation packaging system consisting of psPax2 (Addgene 12260), and pMD2.G (Addgene 12259) using Lipofectamine LTX with plus reagent (ThermoFisher 15338-030). 293FT cells were seeded on gelatin coated 10cm culture dishes and grew until 80% confluent. On the day of transfection, 293FT medium was replaced by fresh medium. To prepare the transfection mix, 5.4 μg of a lentiviral vector, 5.4 μg of psPax2 (Addgene 12260), 1.2 μg of pMD2.G (Addgene 12259) and 12 μl of PLUS reagent were added to 3 ml of Opti-MEM Reduced Serum Media (ThermoFisher 31985-062) and incubated for 5 minutes. After addition of 36 μl of LTX reagent the mixture was incubated for another 30 minutes. The transfection mixture was added dropwise on top of 293FT cells and incubated at 37°C. After 48 hours, the medium with viral particles was collected and fresh medium was added on top of cells. After 24 hours, virus particles were harvested for the second time with medium, and added to the first harvest. All of the harvested virus was filtered using 0.45 um SFCA syringe filter (Nallgene, 190-2545), aliquoted and stored at -80ºC.

#### hIPSC infection

hiPSC cells were harvested using accutase (Stemcell technologies, 07920) and resuspended in mTeSR-E8 Media (Stemcell technologies, 05990) supplemented with 10μM Rock inhibitor (Ri) Y-27632 (Stemcell technologies, 72304). 0.5M cells were aliquoted and mixed with 200μl virus suspension and seeded on one well of 6 well-plate (in replicate). After 24 hours medium with Rock inhibitor was removed and fresh medium without Ri was added on top of cells. After two days, the medium was changed with mTeSR-E8 medium supplemented with 10 μg/ml Blasticidin (TOKU-E, B001), and selection was continued for 10 days.

#### K562 infection

2×10^6^ K562 cells were aliquoted into 10 ml supplemented RPMI medium in duplicate. 100 μl of virus suspension and Polybrene (hexadimethrine bromide, Sigma) at a final concentration of 8 μg/ml were added, and the cell suspensions were centrifuged at 1000g for 30 minutes, resuspended, and seeded in 75 ml culture flasks with additional 15 ml top up with supplemented RPMI media. After 48 hours, cells were passaged into media supplemented with 15 μg/ml Blasticidin (TOKU-E, B001), and selection was continued for 10 days.

#### PiggyBac transposition for hiPSCs

hiPSC were transfected with pB-CAGGS-dCas9-KRAB-MeCP2 or pPB-dCas9-KRAB vectors together with mPBase transposase vector using TransIT®-LT1 Transfection Reagent (Mirus Bio,MIR2300) as recommended by the producer. In summary, cells were harvested with accutase, and resuspended in mTeSR-E8 media supplemented with 10μM Rock inhibitor (Ri) Y-27632 (Stemcell technologies, 72304). Prepare 0.5M cell/ml cell suspension in mTeSR+E8 +Ri media. 400 μl of Opti-MEM Reduced Serum Media (ThermoFisher 31985-062) was aliquoted in a tube and 4μg plasmid DNA (1.8 μg delivery vector, 1.8 μg mPBase, 0.4μg pCS2-GFP) was added. The transfection mixture was incubated at room temperature for 20 minutes, and added onto one well of 6-well plate (Corning, 3516) with 0.5ml of mTeSR-E8+Ri media. After incubating the transfection mix on the plate for another 5 minutes, 1ml of 0.5×10^6^ cell/ml cell suspension was added on top. After 24 hours, the medium was changed to remove Rock inhibitor, and after 48 hours, the medium was changed to mTeSR-E8 supplemented with 10 μg/ml Blasticidin (TOKU-E, B001). Antibiotic selection continued for 10 days.

#### PiggyBac transposition for K562 cells

Cells were transfected with pB-CAGGS-dCas9-KRAB-MeCP2 or pPB-dCas9-KRAB vectors together with mPBase transposase vector using Lipofectamine LTX reagent as described above for 293FT cells. pCS2-GFP vector was used as transfection control and the ratio of delivery vector: mPBas:pCS2-GFP was 1:1:0.2. Blasticidin selection started after 2 days and continued for 10 days.

#### Producing monoclonal hiPSC lines

After 10 days of blasticidin selection, polyclonal cell populations were collected with accutase and seeded 1000 cells into 6-cm dishes precoated with Synthemax™ II-SC Substrate (Corning, 3535) at a concentration of 5 μg/cm2 in mTeSR-E8 media supplemented with 10X CloneR (Stem Cells, 05888). Medium was changed every day, with blasticidin supplemented mTeSR+Plus media (10μg/m). After 10 days, visible colonies were manually transferred into 12 well-plates precoated with Vitronectin XF. Surviving monoclonal cell lines were expanded and used in later analyses.

#### Producing monoclonal K562 lines

After 10 days of blasticidin selection, monoclonal lines were produced by serial dilution as follows: polyclonal cells were counted and serial diluted to an estimated 50 cells in 20ml K562-media and 200μl of suspension was distributed into each well of a 96-well plate. In the first iteration, there was cell growth in 6 wells. Cells from these 6 wells were separately collected, diluted and redistributed into 6 different 96-well plates, and one well from each plate was picked as monoclonal lines.

### Cloning the reporter system

To measure CRISPRi efficacy, we devised a reporter system expressing BFP, GFP, and a guide RNA targeting GFP promoter from the same transient expression vector (Figure 1A). First, the pCS2-GFP vector was linearised with HindIII, and a gBlock (#639) with guide RNA target site was cloned into the GFP promoter region with Gibson assembly (Gibson et al., 2009) to obtain pCS2_CMV_gRNAtarget_GFP vector as a backbone. Meanwhile, the pU6-gRNA5 vector was linearised with BbsI, and oligos with target (#641) and mock (#640) gRNA were cloned after the U6 promoter with Gibson assembly. The U6+gRNA block was amplified from this vector (primers #642 and #643) and cloned into the SapI digested pCS2-BFP vector. The U6+gRNA+BFP block was then amplified from this vector (primers #646 and #647), the PGK promoter was amplified from pKLV2-U6gRNA5(gGFP5)-PGKBFPGFP-W vector (primers #644 and #645), and both cloned into the 5.2kb backbone of pCS2>CMV-GFP-SV40pA><CMV-mRuby2-bGHpA< vector linearised with EcoRi+MluI with Gibson assembly to produce pCS2-iREP-GFP-PGK-BFP-U6-gRNA-iRep or pCS2-iREP-GFP-PGK-BFP-U6-gRNA-mock vectors (all in one reporter system). All Gibson assembly reactions were conducted using NEB Gibson assembly master mix (E2611L) according to the manufacturer’s protocol. Assembled constructs were purified using Monarch PCR purification kit (NEB, T1030S) and used to electroporate into electrocompetent bacteria (NEB, C3020K) according to the manufacturer’s instructions.

### Testing CRISPRi efficacy

Monoclonal and polyclonal hIPSC-dCas9 lines were transfected with pCS2-iREP-GFP-PGK-BFP-U6-gRNA-irep or pCS2-iREP-GFP-PGK-BFP-U6-gRNA-mock vectors as replicates in 12 well plates using TransIT®-LT1 Transfection Reagent (Mirus Bio, MIR2300) according to manufacturer’s instructions and as detailed above. A control transfection for fluorescence gating was conducted using a non-fluorescent plasmid of the same size. Similarly, monoclonal and polyclonal K562-dCas9 lines were transfected with pCS2-iREP-GFP-PGK-BFP-U6-gRNA-irep or pCS2-iREP-GFP-PGK-BFP-U6-gRNA-mock vectors in replicate with Lipofectamine LTX reagent as detailed above, with non-fluorescent plasmid as negative control. Three days after transfection, hIPSC cells were harvested with accutase and K562 cells were collected with centrifugation. All cells were washed and resuspended in PBS+FBS (2%). Harvested cells were analyzed in CytoFLEX flow cytometer and FlowJo analysis software (Beckman Coulter). CRISPRi efficacy was quantified as percent activity - the percent decrease in median GFP level in BFP positive cells.

### Generating the CRISPRi tiling library

#### Design

To determine the optimal target window of dCas9-KRAB-MeCP2 in iPSCs, three sub-libraries were designed (Supplementary Figure 1). First, the TSS tiling library placed guides relative to the transcription start sites in human iPSCs, as identified from the CAGE annotations in the Fantom database (Lizio et al., 2015) according to the GRCh38 reference genome. Within the TSS tiling library, the “TSS Tiling, Extra” library tiled from 0 to +100bp of TSS site for the single transcript of 882 genes essential in human iPSCs (Figure 2A); the “TSS Tiling, Different” library tiled from -200 to +300 bp of TSS site for the top two transcripts of 20 essential genes for which the TSS annotation in human iPSCs did not match the canonical one (Figure 2B); and the “TSS Tiling, Other” library tiled top two transcripts of 20 essential genes. Second, the Gene tiling library targeted guides to all protospacer adjacent motifs in coding sequence of a single transcript of 451 Hart essential genes, and 36 non-essential genes (Figure 2C). Finally, 200 non-targeting guides were used as control (Supplementary Table 1).

#### Cloning and titration

sgRNAs were synthesized as complex oligonucleotide pools with several sub-pools (Genscript). Subpools were amplified with PCR, and Gibson assembly homology sequences were added to sgRNA sequences in a second round of amplification (primers #745 and #746). The lentiviral backbone vector pKLV2-U6gRNA5(BbsI)-ccdb-PGKpuroBFP-W (AddGene: 67974) (Tzelepis et al., 2016) was linearised with BbsI. Amplified sgRNA pools were inserted into the vector backbone with Gibson assembly reaction using NEB Gibson assembly master mix (E2611L) according to the manufacturer’s protocol. Assembled constructs were purified using Monarch PCR purification kit (NEB, T1030S). The assembled plasmid library was electroporated into electrocompetent bacteria (NEB, C3020K) according to the manufacturer’s instructions as 3 reactions. 5μl of recovered bacteria was diluted 1:10 three times, and dilutions were seeded on ampicillin plates, while the remaining bacteria were seeded in liquid culture with ampicillin selection (100 μg/ml). The next day, bacterial colonies on agar plates were counted to calculate the coverage of the library, which was between 80x and 100x. The plasmid library pool was isolated from overnight bacteria culture using QIAGEN HiSpeed Plasmid Midi (small pooled library) of maxi (TSS tiling library) kits (QIAGEN, 12643 and 12662). Lentivirus was produced from cloned plasmid pools in 293FT cells as described above. To determine virus titer hiPSC-(Fiaj-1) dCas9-KRAB-MeCP2 cells were suspended as 0.16M cells/ml suspension. 5 × 2.5 ml of cell suspensions were mixed with five different amounts of virus between 20μl to 100μl. 1ml of each virus cell mixture was seeded one well of vitronectin pre-coated 12 well plate. After 3 days, cells were collected with accutase, washed, and resuspended in FACS buffer (PBS with 2% FBS) and analysed by FACS (Cytoflex, BD). Best fit line formula of percentage BFP positive cells against virus volume was used to calculate the volume of virus required for 0.3 multiplicity of infection (MOI).

### Reamplification and titration of the Dolcetto CRISPRi library

Dolcetto library (Sanson et al., 2018) was acquired from Addgene (#1000000114) as two plasmid pools, each with a half-library (Set A and B) in the XPR_500 backbone. Only Dolcetto Set A plasmid pool was used in this study. 350 ng Dolcetto setA library pool was used to transform 100 μl electrocompetent bacteria (NEB, C3020K), split into 4 × 25 μl electroporations. Electroporated bacteria were recovered in 3ml recovery media and all 4 cultures were mixed together. 5 ul of the culture mix was diluted 1:10 serial diluted 5 times and each dilution seeded on LB agar+ Amp plates. Remaining transformed bacteria culture seeded in 0.5ml LB+ Ampicillin (100 μg/ml) was left overnight. Plasmid library pool was isolated from overnight culture using HiSpeed maxi plasmid isolation kit (QIAGEN, 12662). Lentivirus was produced from the amplified plasmid pool as described above. To titrate the virus for hiPSCs, hIPSC+dCas9_KRAB+MeCP2 lines were collected with accutase and 1 M cells were resuspended in mTeSR-Plus +Ri media at 1.3×10^5^ cells/ml. Different volumes of virus prep from 0 μl to 1.6μl were mixed with mTeSR-Plus +Ri media up to 600 μl. 600 μl of cell suspension was added on top of each virus dilution and mixed. 50 μl of each virus cell mixture were seeded into 10 wells of each of 96-well plates, resulting in two plates with 6 different virus/cell ratios. On day 2 post-infection, the medium in one plate was changed with mTeSR-Plus, and in the other plate with mTeSR-Plus + Puromycin (0.5 μg/ml). On day 4 post-infection, the medium in each well was replaced by 100μl mTeSR-Plus medium + 20μl MTS dye (Promega, G3582) to compare living cell amounts between plates. The plates were incubated at 37 C for 2 hours, followed by the addition of 25μl of 10% SDS solution on each well. Plates were analyzed in Multiskan™ GO Microplate Spectrophotometer plate reader (Thermo Scientific, 51119200) at wavelengths of 490 nm and 700 nm. Raw reads were adjusted after blank and background (700 nm) readings were subtracted from target readings (490 nm). Virus titer was determined from the ratio of puromycin treated and untreated cells. To titrate the virus for K562 cells, the cells were collected, counted, and aliquoted to have 50,000 cells per well, 12 wells for each of 6 virus titers. Different amounts of virus prep were mixed with cell suspensions and distributed on a 96-well plate. Each well was topped up with supplemented RPMI media + Polybrene (8ug/ml final concentration) up to 150 ul. Plates were centrifuged at 1000 rcf for 30 minutes and resuspended. On day 3 after infection, cells were split into two 96-well plates. Cells in one plate were resuspended in supplemented RPMI media and the other plate in supplemented RPMI + puromycin (2μg/ml) media. On day 6 after the infection relative amounts of cells in each well were determined by MTS assay as described above.

### CRISPRi screening

#### Human iPSCs

6×10^7^ hiPS cells were mixed with lentiviral Dolcetto library to achieve estimated 0.3 MOI and seeded on 2x Vitronectin coated 5-layer flasks at 36K cells/cm2 density in mTeSR-Plus medium with Rock Inhibitor (Ri, 10μM, Y-27632, Stemcell technologies, 72304). The next day, the medium was changed to remove Ri. On day 3 post infection, puromycin selection was started at 0.5 ug/ml, and continued through the screen duration (17 to 22 days). Cells were passaged at 90% confluency and seeded back at 16K cells/cm2 density for the rest of the screen. The remaining cell pellets were aliquoted and frozen for genomic DNA extraction.

#### K562 cells

75×10^6^ cells were resuspended in supplemented RPMI+ polybrene (8μg/ml) media and mixed with Dolcetto library virus aiming for MOI of 0.3. The cell+virus suspension was centrifuged at 1000 rcf for 30 minutes. The cells were then resuspended in the same media and plated into two T150cm2 plates for a final concentration of 0.45×10^6^ cells/ml. The infected cells were selected with puromycin (2ug/ml) starting from day 1 after infection until day 8. After complete selection, the cells in culture were passaged every 3 or 4 days, and plated back at 0.1×10^6^ cells/ml density. Aliquoted pellets of at least 40×10^6^ cells were frozen for genomic DNA extraction.

#### Genomic DNA isolation and sequencing library preparation

Aliqouted cell pellets were thawed at room temperature and resuspended in 100 mM Tris-HCl, pH 8.0, 5 mM EDTA, 200 mM NaCl, 0.2% SDS and 1 mg/ml Proteinase K. After overnight incubation at 55oC, RNase was added on top of the cell suspension to the final concentration of 10 μg/μl, followed by 3h incubation at 37oC. Genomic DNA was precipitated with 100% isopropanol, spooled out, washed in 70% EtOH and air dried at room temperature. Following resuspension in TE buffer overnight, DNA was quantified with Quant-iT Broad Range kit (Q33130, ThermoFisher). gRNA cassettes were amplified from genomic DNA with two consecutive PCR reactions taking the target library coverage into account (used genomic DNA corresponding cell number for 500x for final timepoints, minimum 250x for interval timepoints). The first reaction with Q5 Hot Start High-Fidelity 2X Master Mix (NEB) amplifies the gRNA cassette (primers #1 and #2 for CRISPRi and small pooled screen libraries, primers #1 and #638 for Dolcetto library). PCR reactions were purified with QIAquick PCR Purification Kit (Qiagen, 28106) and quantified with nanodrop. Each library was diluted to 1ng/μl and used as template for the second PCR reaction where sequencing adaptors with index sequences were added as described before (primers #15 and #NN indexing) (Tzelepis et al. 2016). PCR reactions were purified with 0.7X AMPure XP beads (Agencourt AMPure XP beads; Beckman, Cat.no. A63881), quantified with Quant-it High Sensitivity Kit (Q33120, ThermoFisher), pooled and single end sequenced using primer #16.

### Arrayed gRNA cloning, lentivirus production

#### Cloning

Each guide RNA was ordered as top and bottom strand oligos mixed in single well creating double stranded DNA with cloning overhangs (Sigma Aldrich, Supplementary Table 4). Oligos were phosphorylated with T4 Polynucleotide Kinase (NEB M0201) at 37oC for 30 minutes. pKLV2-U6gRNA5(BbsI)-PGKpuro2AmCherry-W (Addgene 67977) (Tzelepis et al. 2016) was linearised with BbsI at 37oC overnight. Phosphorylated gRNA oligos were cloned into linearized backbone with T4 DNA ligase (NEB M0202*) by incubating 1h at room temperature. 3 μl of the ligation products were used to transform 20μl of zymo mix&go 10B competent cells (Zymo Research, T3020). Transformed cells were grown in 2ml 2XLB media overnight. gRNA plasmids were isolated using QIAprep Turbo Miniprep Kit (Qiagen, 27191). gRNA sequences were confirmed by Sanger sequencing.

#### Lentivirus production

To produce arrayed gRNA lentivirus, the flat bottom 96 well plates were coated with 0.1% gelatin, and 293T cells were seeded as 20K cells/well. In day 1, for each well, 0.1ug LV transfer plasmid, 0.1ug psPAX2, and 0.02 ug pMD2.G were used to transfect 293T cells with Lipofectamine LTX Reagent (15338100, Thermo Fisher) by scaling manufacturer’s protocol. Cell medium with the virus were collected from each well after 48 hours, centrifuged at 500 rcf for 20 minutes, and supernatant was collected carefully without disturbing the pellet.

### Cell phenotyping

#### Growth assays with live imaging system

Cell growth was measured in Incucyte S3 Live Imaging System with confluence as a measure of cell number in different assays. Additionally, to validate confluence vs cell number correlation, cells were seeded in a 96 well plate at different densities and cell growth was measured by taking images every two hours with a 10X objective and 5 frames per well for 24h. After final measurement of confluence in Incucyte live imaging system for each, we compared the optical density (OD) value obtained to the MTS assay and found it to be highly correlated (Supplementary Figure 10).

Comparison of WT and ARID1A^+/-^ lines. Wild type and ARID1A^+/-^ Kolf hiPSC lines were seeded at 20,000/cm2 density in 12-well plates (6 well each) in mTeSR Plus medium supplemented with Rock inhibitor (Ri). The next day, medium was changed to remove Ri and the plate was placed in Incucyte S3 live imaging system (Sartorius) and imaged every 2 hours for 50 hours using 10x objective taking 16 frames per well. The growth rate was measured as percent confluency and normalized to the first time point.

#### Oprozomib treatment measurements

Wild type and ARID1A^+/-^ Kolf hiPSC lines were seeded at 20,000/cm2 and 30,000/cm2 density respectively in 12 wells of 24 well plate. After 36 hours, both lines were at approximately 25% confluence. At this point, medium was replaced with mTeSR-Plus supplemented with different concentrations of Oprozomib (APExBIO, A1937) (10uM, 2uM, 0.4uM, 0.08uM, 0,02uM). The plate was then placed in the Incucyte live imaging system, and imaged every 2 hours for 26 hours using 10x objective taking 16 frames per well. At each time point, the growth rate was measured as percent confluency, and normalised to the first time point.

#### CRISPR vs CRISPRi comparison

Monoclonal Fiaj-1_dCas9-KRAB-MeCP and Fiaj-1_Cas9 lines were collected, counted and infected with targeting and non-targeting guides (Supplementary Table 3). Infected cells were seeded on two wells of 24 plates and monitored in Incucyte S3 Live Cell imaging System with 10x objective taking 16 frames per well every two hours. Cell death was determined as decrease in confluency after media change.

#### Cell staining and FACS

Wild type and ARID1A^+/-^ lines were infected with lentivirus with EZH2 targeting guides and seeded in 12 well plates. Infected cells were selected with puromycin for 9 days. Cells were collected with accutase and counted. 5×10^6^ cells were aliquoted and washed once with FACS buffer (5% FBS in PBS). Cells were fixed in 1% PFA for 30 minutes at room temperature and blocked in FACS buffer for another 30 minutes. 1×10^6^ cells were aliquoted in tubes and incubated with stem cell marker, perCP-CY-5.5-Mouse antihuman-TRA-1-60 antibody (BD Pharmingen, 561573) for 1 hour at room temperature. HEK293T cells were used as a negative control for antibody sensitivity. Stained cells were analyzed in CytoFLEX flow cytometer and with FlowJo analysis software (Beckman Coulter).

### Data analysis

gRNA sequences were counted from fasta files and aligned to corresponding library maps without allowing any mismatches. Uniformity of plasmid libraries was determined using the Gini index as implemented by Wang *et al*. (Wang et al., 2019). Average sequencing coverage of all libraries was around 400x. When all technical replicates were combined together, a single screen had around 2000x coverage for final time points. Raw counts in each sequencing sample were normalized and transformed to log_2_ scale according to the formula: log_2_(((reads per guide/sum of all read counts)*10^6^)+1). Log_2_ fold change per guide was calculated by subtracting log_2_ transformed and normalised read counts of final time points from plasmid library sequences. To determine essential and non-essential genes, Bagel v2 (Kim and Hart, 2021) was used with default parameters. To find differentially essential genes between ARID1A^+/+^ and ARID1A ^+/-^ lines, we calculated the mean and standard deviation of the difference across all genes, and assigned the genes with values of at least two standard deviations away from the mean to be differentially essential.

## SUPPLEMENTARY DATA

are available with the paper.

**Supplementary Table 1:** CRISPRi Tiling library guide RNA and log fold change data

**Supplementary Table 2:** Data for pooled screen:

Guide RNA read counts for whole genome screens in iPSCs

Gene mean log_2_ fold changes for whole genome screens in iPSCs

Gene Bayes factor values for whole genome screens in iPSCs

Area Under ROC Curve values for whole genome screens in iPSCs

Whole genome library plasmid quality control data

Gene level summaries of CRISPR and CRISPRi screens Guide RNA read counts for followup pooled library

Gene Bayes factor values for followup pooled library

**Supplementary Table 3:** Guide RNAs used in arrayed screens

**Supplementary Table 4:** Primer sequences

## Supplementary Note

Comparison of our screens in ARID1A^+/-^ lines with screens in the DepMap database.

## ACKNOWLEDGEMENTS

SU was supported by Janet Thornton Fellowship. PH, LC, YZ, KU, CD, JS, GN, OEG, SG, BN, OMD, LP were supported by Wellcome (206194). KA was supported by the Estonian Research council (PSG415) and the Archimedes Foundation (EXCITE TK148).

