## SUPPLEMENTARY DATA for "Optimised whole-genome CRISPR interference screens identify ARID1A-dependent growth regulators in human induced pluripotent stem cells"

**SUPPLEMENTARY FIGURES**

**Supplementary Figure 1.**

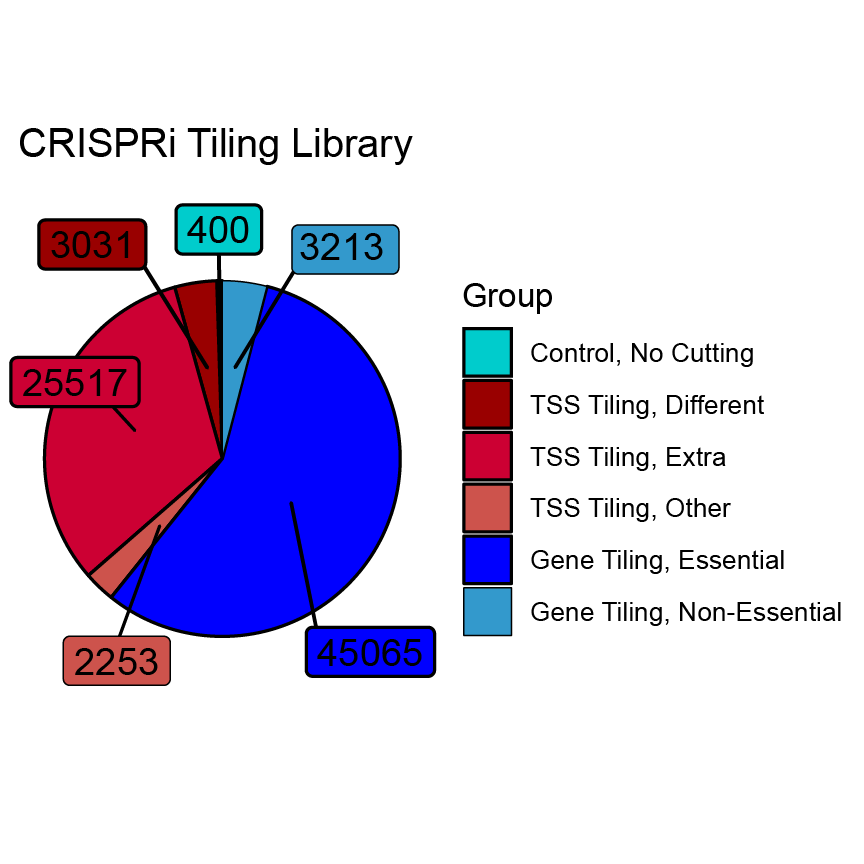

**Supplementary Figure 1**: Composition of CRISPRi tiling guide RNA library used in Figure 2. Numbers in boxes: guide RNA numbers in each sub-library. Teal: control library with no genomic targets. Red shades: transcription start site tiling library (“TSS Tiling, Extra”) tiling from 0 to +100bp of TSS site for the single transcript of 882 genes essential in human iPSCs; different TSS tiling library (“TSS Tiling, Different”) tiling -200 to +300 bp of TSS site for the top two transcripts of 20 essential genes for which the TSS annotation in human iPSCs did not match the canonical one; other transcript tiling library (“TSS Tiling, Other”) tiling top two transcripts of 20 essential genes. Blue shades: gene tiling library targeting all protospacer adjacent motifs in coding sequence of a single transcript of 451 Hart essential genes (blue; Hart et al., 2014) and 36 non-essential genes (teal). Contents of the libraries are provided in Supplementary Table 1.

**Supplementary Figure 2**.

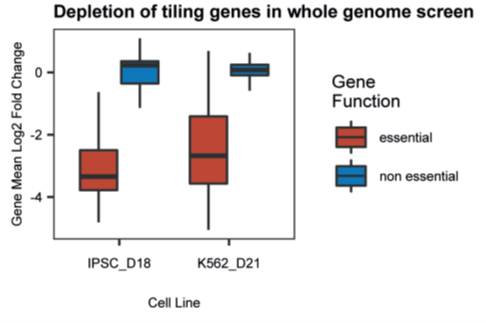

**Supplementary Figure 2**: Gene-mean log2-fold change (y-axis) of genes in the transcription start site tiling library (Figure 2) in whole genome screen in iPSCs and K562 lines (x-axis). Box: median and quartiles; whiskers: 95th percentile; red: core essential genes, blue: non-essential genes.

**Supplementary Figure 3.**

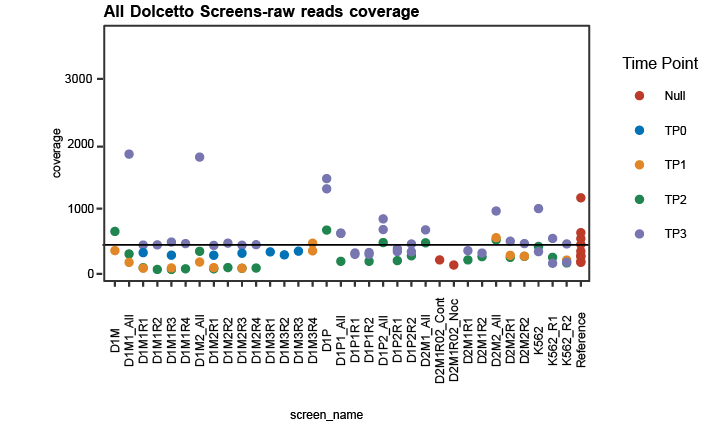

**Supplementary Figure 3**: Sequencing coverage (y-axis) of sequencing libraries (x-axis) of whole genome screens conducted using Dolcetto library (56,554 guide RNAs in total). Screen names; D: Donor, M: Monoclone, P: Polyclone, R: Replicate. Red: reference libraries, blue: timepoint 0 (days 3-5), yellow: timepoint 1 (days 9-11), green: timepoint 2 (days 13-15), purple: timepoint 3 (days 18-22), black line: average screen coverage (445X).

**Supplementary Figure 4**.

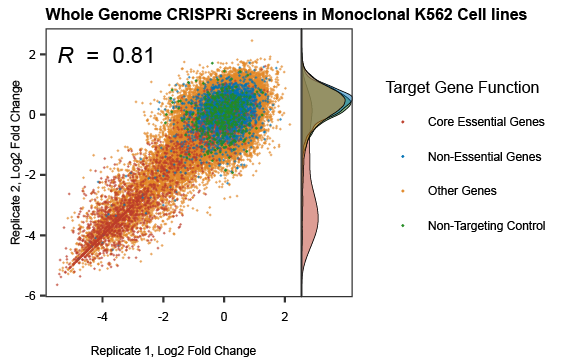

**Supplementary Figure 4**: Reproducibility of genome-wide screens in monoclonal K562 lines. gRNA log2-fold change in replicate 1 (y-axis) and replicate 2 (x-axis). Red: essential genes; blue: non-essential genes; yellow: other genes; green: non-targeting controls.

**Supplementary Figure 5.**

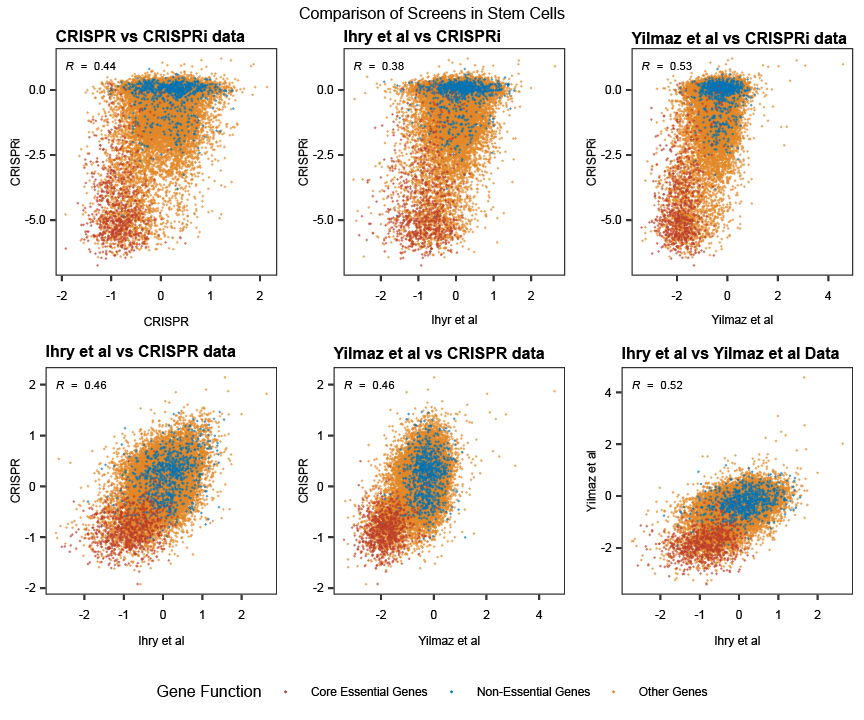

**Supplementary Figure 5**: Concordance of gene average log2-fold changes (x- and y-axes) of in-house CRISPR and CRISPRi screens in hiPSC with previously published CRISPR screens in diploid (Ihry *et al.*) and haploid (Yilmaz *et al.*) human embryonic stem cells (hESC). Red: essential genes; blue: non-essential genes; yellow: other genes. R: Pearson’s correlation coefficient.

**Supplementary Figure 6**.

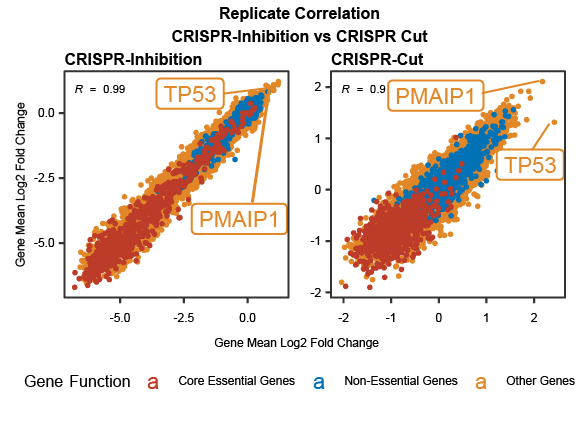

**Supplementary Figure 6**: Reproducibility and p53 effect in whole genome CRISPRi (left) and CRISPR (right) screens in monoclonal hiPSC lines. Gene mean log2 fold change value for replicate 1 (x-axis) and replicate 2 (y-axis). Data points for p53 and PMAIP1 genes are shown with an arrow. Red: essential genes; blue: non-essential genes; yellow: other genes.

**Supplementary Figure 7**.

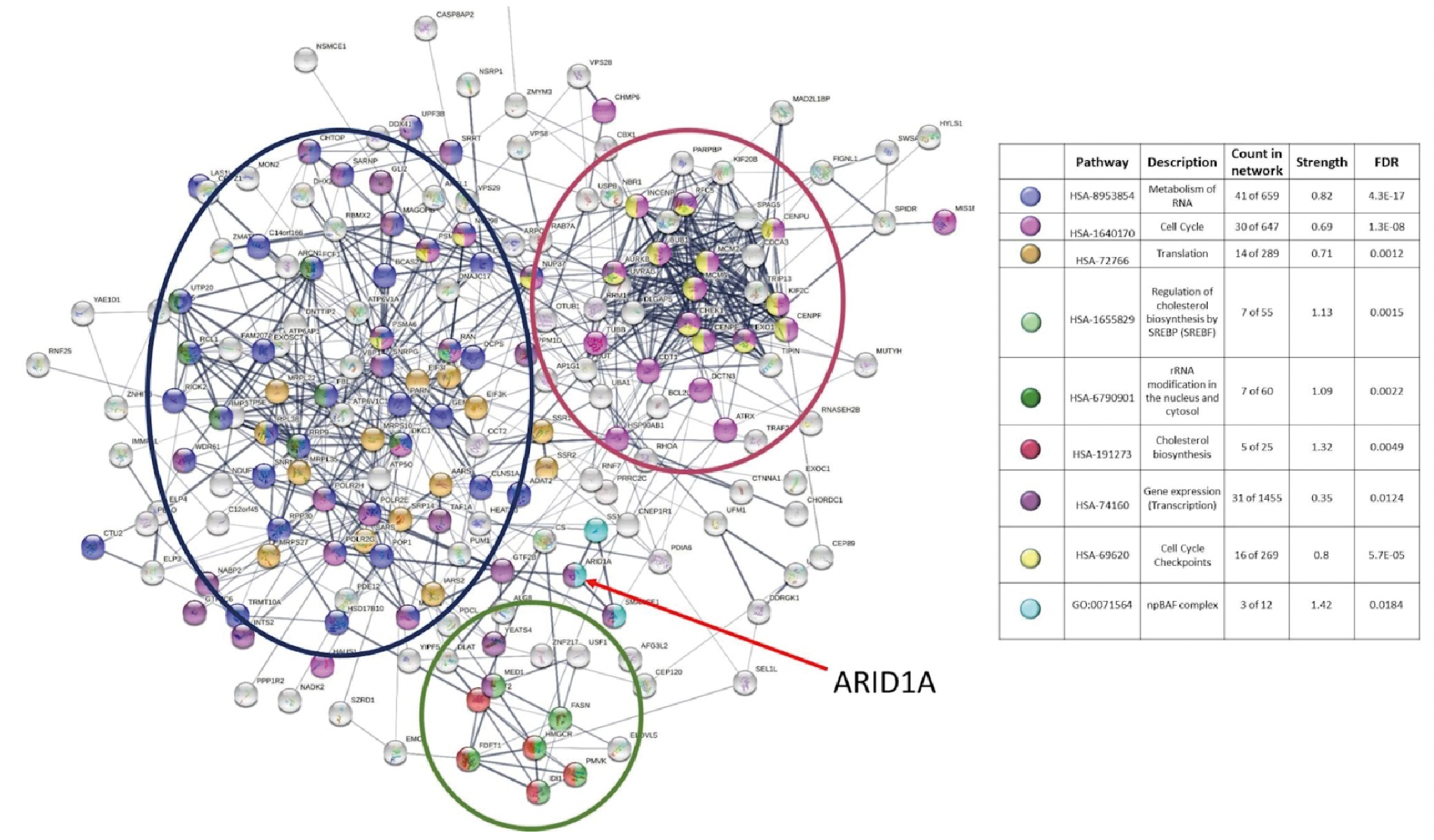

**Supplementary Figure 7.** STRING analysis of ARID1A. Large ellipses: manually curated clusters.

**Supplementary Figure 8**.

Non-targeting guide RNA	EZH2-targeting guide RNA

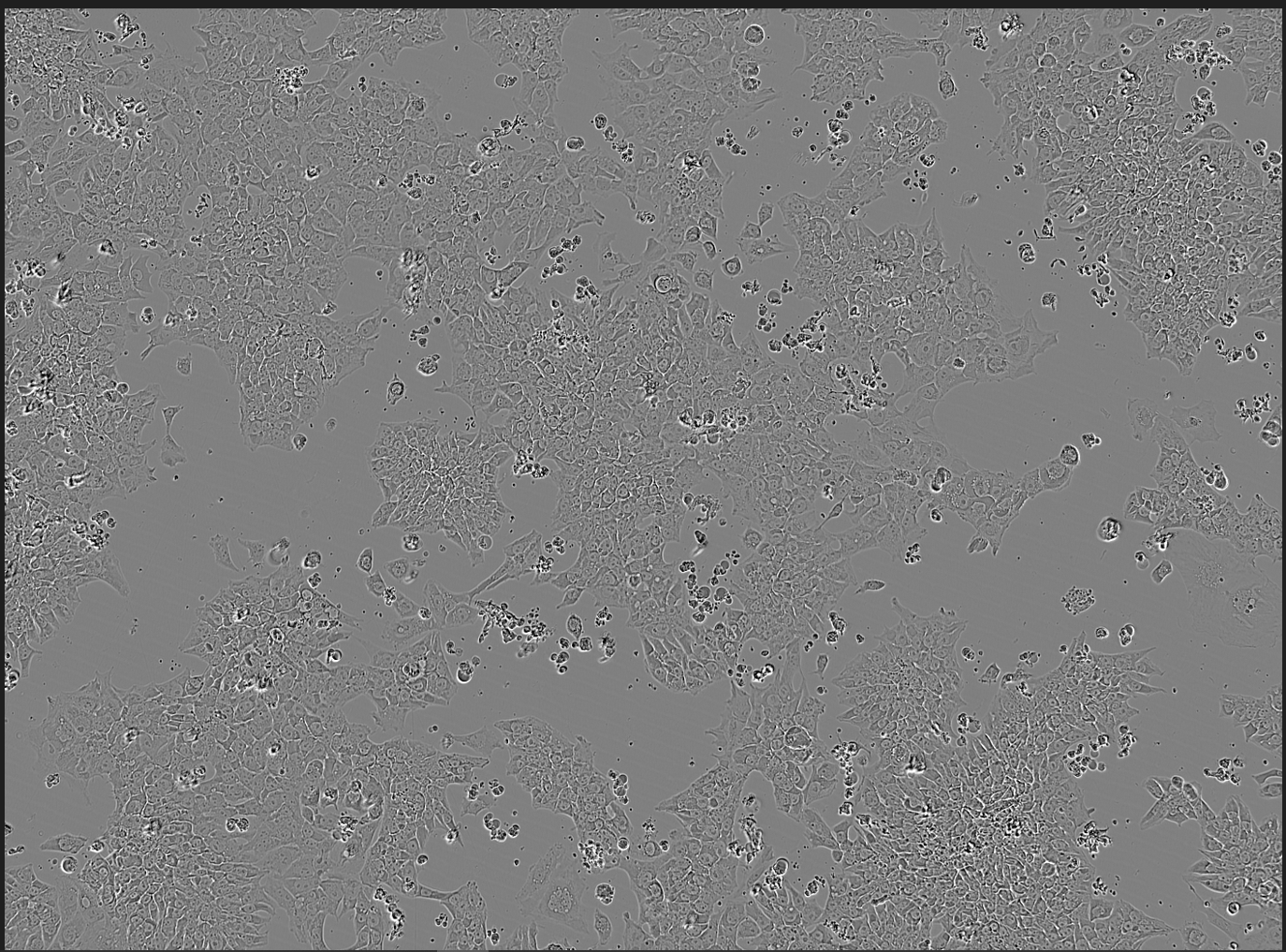

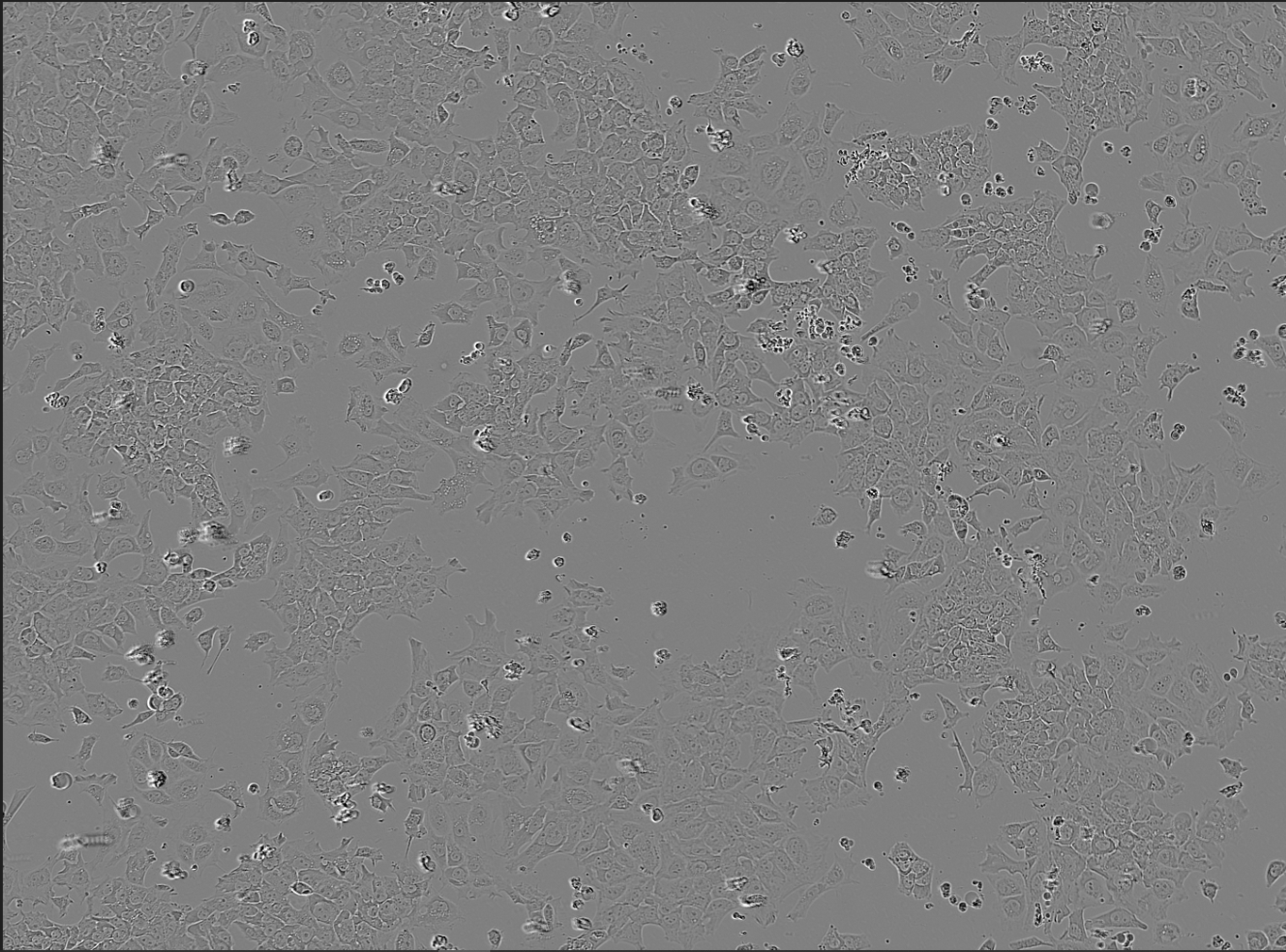

**Supplementary Figure 8.** Microscope images from Incucyte live cell imaging system (10X objective) of ARID1A^+/+^ hiPSCs infected with non-targeting guide RNA (left) and a guide RNA targeting EZH2 gene (right).

**Supplementary Figure 9.**

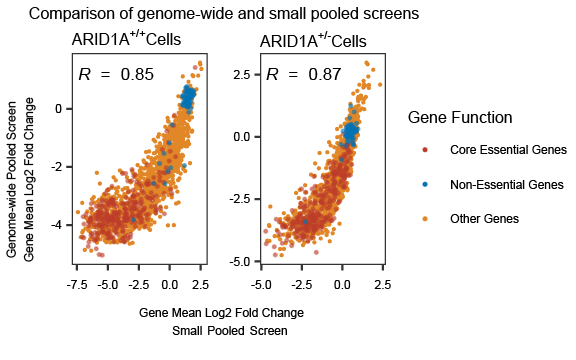

**Supplementary Figure 9.** Reproducibility of gene average log2-fold change (markers) in genome-wide (y-axis) and small-pooled screens (x-axis) in ARID1A^+/+^ (left panel) and ARID1A^+/-^ (right panel) hiPSC lines. Red: essential genes; blue: non-essential genes; yellow: other genes.

**Supplementary Figure 10**.

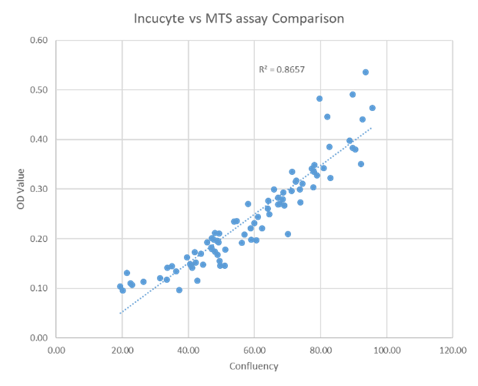

**Supplementary Figure 10.** Concordance between confluency of hiPSC under the microscope (x-axis) and cell number measured as OD value by MTS assay (y-axis) for each well of a 96 well plate. Diagonal line: linear model fit.

**Supplementary Figure 11.**

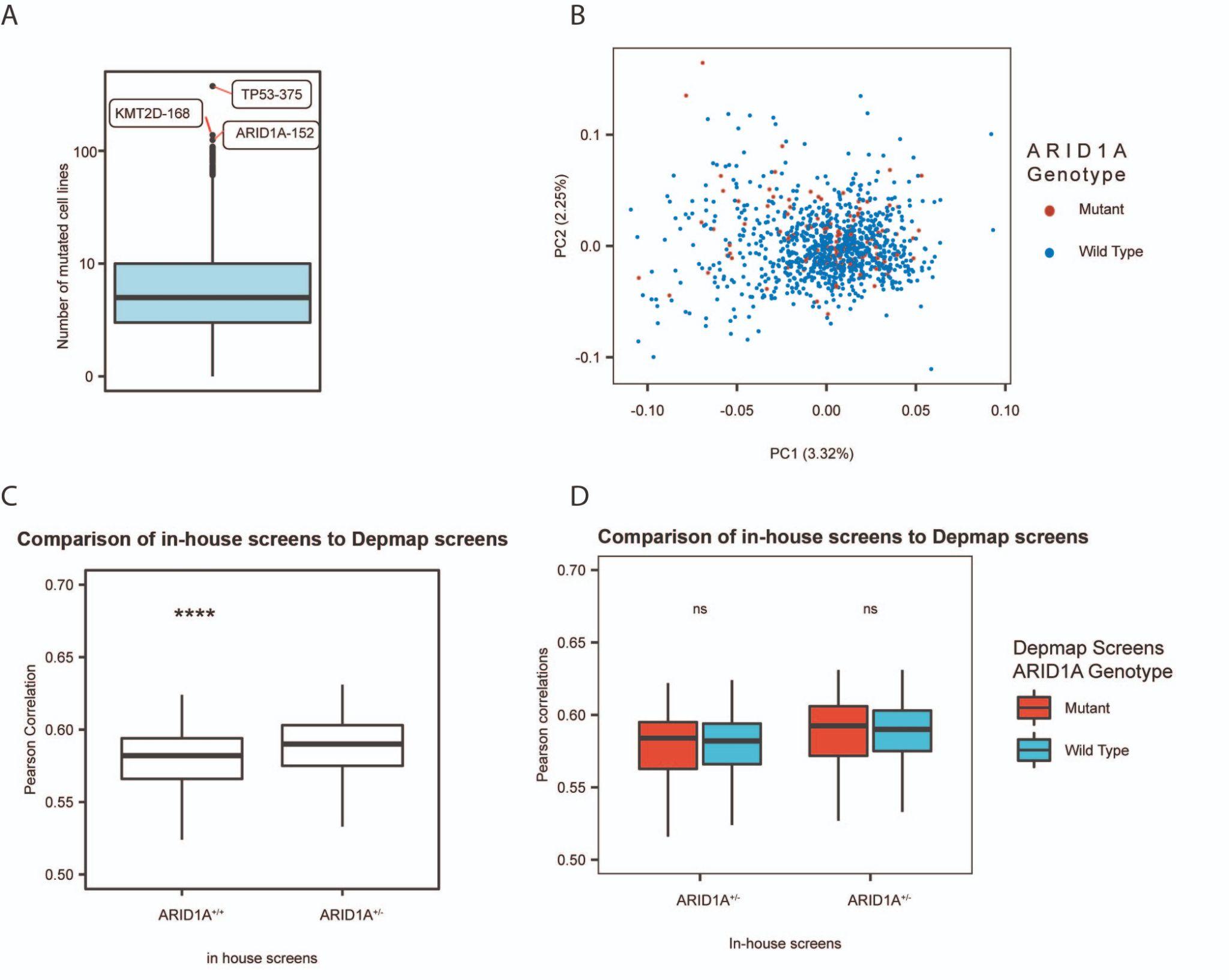

**Supplementary Figure 11**: Comparison of in-house CRISPRi screens in ARID1A+/- and ARID1a+/+ hiPSC lines to DepMap screens. **A.** Number of mutated cell lines (y-axis) per gene. Labels: Gene and number of unique cancer lines with at least one mutation in a particular gene. Box: median and quartiles; whiskers: 95th percentile. **B.** PCA plot of DepMap CRISPR Gene Effect scores labeled according to ARID1A genotype of the cell line. Red: ARID1A mutant lines, Blue: ARID1A wild type lines. **C**. Correlation of in-house CRISPRi screens’ Bayes Factor values to DepMap CRISPR Gene Effect scores. ARID1A^+/+^: CRISPRi screen in ARID1A^+/+^ hiPSC line, ARID1A^+/-^: CRISPRi screen in ARID1A^+/-^ hiPSC line. **D.** Correlation of in-house CRISPRi screens’ Bayes Factor values to DepMap CRISPR Gene Effect scores separated by ARID1A genotype of DepMap cancer lines; Red: DepMap cancer lines with a mutation in ARID1A Gene. Blue: DepMap Cancer lines without ARID1A mutation. ARID1A^+/+^: CRISPRi screen in ARID1A^+/+^ hiPSC line, ARID1A^+/-^: CRISPRi screen in ARID1A^+/-^ hiPSC line.

**SUPPLEMENTARY FIGURE LEGENDS**

**Supplementary Figure 1**: Composition of CRISPRi tiling guide RNA library used in Figure 2. Numbers in boxes: guide RNA numbers in each sub-library. Teal: control library with no genomic targets. Red shades: transcription start site tiling library (“TSS Tiling, Extra”) tiling from 0 to +100bp of TSS site for the single transcript of 882 genes essential in human iPSCs; different TSS tiling library (“TSS Tiling, Different”) tiling -200 to +300 bp of TSS site for the top two transcripts of 20 essential genes for which the TSS annotation in human iPSCs did not match the canonical one; other transcript tiling library (“TSS Tiling, Other”) tiling top two transcripts of 20 essential genes. Blue shades: gene tiling library targeting all protospacer adjacent motifs in coding sequence of a single transcript of 451 Hart essential genes (blue; Hart et al., 2014) and 36 non-essential genes (teal). Contents of the libraries are provided in Supplementary Table 1.

**Supplementary Figure 2**: Gene-mean log2-fold change (y-axis) of genes in the transcription start site tiling library (Figure 2) in whole genome screen in iPSCs and K562 lines (x-axis). Box: median and quartiles; whiskers: 95th percentile; red: core essential genes, blue: non-essential genes.

**Supplementary Figure 3**: Sequencing coverage (y-axis) of sequencing libraries (x-axis) of whole genome screens conducted using Dolcetto library (56,554 guide RNAs in total). Screen names; D: Donor, M: Monoclone, P: Polyclone, R: Replicate. Red: reference libraries, blue: timepoint 0 (days 3-5), yellow: timepoint 1 (days 9-11), green: timepoint 2 (days 13-15), purple: timepoint 3 (days 18-22), black line: average screen coverage (445X).

**Supplementary Figure 4**: Reproducibility of genome-wide screens in monoclonal K562 lines. gRNA log2-fold change in replicate 1 (y-axis) and replicate 2 (x-axis). Red: essential genes; blue: non-essential genes; yellow: other genes; green: non-targeting controls.

**Supplementary Figure 5**: Concordance of gene average log2-fold changes (x- and y-axes) of in-house CRISPR and CRISPRi screens in hiPSC with previously published CRISPR screens in diploid (Ihry *et al.*) and haploid (Yilmaz *et al.*) human embryonic stem cells (hESC). Red: essential genes; blue: non-essential genes; yellow: other genes. R: Pearson’s correlation coefficient.

**Supplementary Figure 6**: Reproducibility and p53 effect in whole genome CRISPRi (left) and CRISPR (right) screens in monoclonal hiPSC lines. Gene mean Log2 fold change value for replicate 1 (x-axis) and replicate 2 (y-axis). Data points for p53 and PMAIP1 genes are shown with an arrow. Red: essential genes; blue: non-essential genes; yellow: other genes.

**Supplementary Figure 7.** STRING analysis of ARID1A. Large ellipses: manually curated clusters.

**Supplementary Figure 8**: Microscope images from Incucyte live cell imaging system (10X objective) of ARID1A^+/+^ hiPSCs infected with non-targeting guide RNA (left) and a guide RNA targeting EZH2 gene (right).

**Supplementary Figure 9.** Reproducibility of gene average log2-fold change (markers) in genome-wide (y-axis) and small-pooled screens (x-axis) in ARID1A^+/+^ (left panel) and ARID1A^+/-^ (right panel) hiPSC lines. Red: essential genes; blue: non-essential genes; yellow: other genes.

**Supplementary Figure 10.** Concordance between confluency of hiPSC under the microscope (x-axis) and cell number measured as OD value by MTS assay (y-axis) for each well of a 96 well plate. Diagonal line: linear model fit.

**Supplementary Figure 11**: Comparison of in-house CRISPRi screens in ARID1A+/- and ARID1a+/+ hiPSC lines to DepMap screens. **A.** Number of mutated cell lines (y-axis) per gene. Labels: Gene and number of unique cancer lines with at least one mutation in a particular gene. Box: median and quartiles; whiskers: 95th percentile. **B.** PCA plot of DepMap CRISPR Gene Effect scores labeled according to ARID1A genotype of the cell line. Red: ARID1A mutant lines, Blue: ARID1A wild type lines. **C**. Correlation of in-house CRISPRi screens’ Bayes Factor values to DepMap CRISPR Gene Effect scores. ARID1A^+/+^: CRISPRi screen in ARID1A^+/+^ hiPSC line, ARID1A^+/-^: CRISPRi screen in ARID1A^+/-^ hiPSC line. **D.** Correlation of in-house CRISPRi screens’ Bayes Factor values to DepMap CRISPR Gene Effect scores separated by ARID1A genotype of DepMap cancer lines; Red: DepMap cancer lines with a mutation in ARID1A Gene. Blue: DepMap Cancer lines without ARID1A mutation. ARID1A^+/+^: CRISPRi screen in ARID1A^+/+^ hiPSC line, ARID1A^+/-^: CRISPRi screen in ARID1A^+/-^ hiPSC line.

**Supplementary Note**

Since our results in whole genome screens show gene interactions important for carcinogenesis, we asked if the sensitizing effect of the ARID1A mutation is similar to those of the damaging mutations in the cancer cell lines. To answer this question, we compared the survival of cancer cell lines upon gene perturbation in ARID1A wild-type and mutant contexts using data from the DepMap project (Dempster et al., 2021; Pacini et al., 2021). Among the 1,755 cancer lines in the database, 152 have at least one damaging mutation in the ARID1A gene (data: Q1/2022), making it one of the most mutated genes in this set of cancer lines (Supplementary Figure 11A). However, ARID1A mutation status was not associated with broad changes in gene essentiality in general (Supplementary Figure 11B). We did observe that gene essentiality estimates from our screen in the ARID1A+/- hiPSC line are significantly more correlated to the ones in cancer cell lines compared to the wild-type line (Supplementary Figure 11C). Still, this effect is small, and the correlation is not affected by ARID1A mutation status of the cancer line (Supplementary Figure 11D).
