## Supplementary Table 4 for "Optimised whole-genome CRISPR interference screens identify ARID1A-dependent growth regulators in human induced pluripotent stem cells"

| Primer | Sequence | Name |
| --- | --- | --- |
| #639 | TGCCTGGGGACGTCGGAGCATACCTCATCAGGAACATGTTGGTGATTTAGGTGACACTATAGAATACAAGCT | Gibson-target-iREP |
| #640 | GCTTTATATATCTTGTGGAAAGGACGAAACACCGGTTAAGCGACTTCGGCCAGGTTTAAGAGCTATGCTGGAAACAGCA | Gibson-mock-grna |
| #641 | GCTTTATATATCTTGTGGAAAGGACGAAACACCGTACCTCATCAGGAACATGTGTTTAAGAGCTATGCTGGAAACAGCA | Gibson-iREP-gRNA |
| #642 | GGGCCTCTTCGCTATTACGCCAGACGCGTCCAAGGTCGGGCA | GA-pCS2-U6 |
| #643 | ATACGCCATATTGAATTGGCTATGGTCGACACTAAAGGGAACAAAAGCGGATCC | GA-pCS2-scaffold |
| #644 | CCGCCAGTGTGATGGATATCTGCAGTTAATTTAGCTTGTGCCCCAGTTTGCT | GA-BFP-bGHpA |
| #645 | CTTCCTGCCCGACCTTGGACGCGTAATTCTACCGGGTAGGGGAGGCGC | GA-U6-PGK |
| 745 | GGCTTTATATATCTTGTGGAAAGGACGAAA | ssoligo-PCR-F |
| 746 | ACTTGCTATGCTGTTTCCAGCATAGCTCTT | ssoligo-PCR-R |
| #1 | ACACTCTTTCCCTACACGACGCTCTTCCGATCTCTTGTGGAAAGGACGAAACA | Sequencing_gRNA_library_amplification |
| #2 | TCGGCATTCCTGCTGAACCGCTCTTCCGATCTCTAAAGCGCATGCTCCAGAC | Sequencing_gRNA_library_amplification |
| #638 | TCGGCATTCCTGCTGAACCGCTCTTCCGATCTTCTACTATTCTTTCCCCTGCACTGT | Dolcetto-lib-R |
| #15 | AATGATACGGCGACCACCGAGATCTACACTCTTTCCCTACACGACGCTCTTCCGATCT | Sequencing_indexing_PCR_indexing |
| #NN | CAAGCAGAAGACGGCATACGAGATN11GAGATCGGTCTCGGCATTCCTGCTGAACCGCTCTTCCGATCT | Sequencing_indexing_PCR |
| #16 | TCTTCCGATCTCTTGTGGAAAGGACGAAACACCG | Sequencing |
